# Quorum Sensing Regulates Virulence Factors in the Coral Pathogen *Vibrio coralliilyticus*

**DOI:** 10.1101/2024.06.10.598281

**Authors:** Victoria N. Lydick, Shir Mass, Robert Pepin, Ram Podicheti, Emra Klempic, Douglas B. Rusch, Blake Ushijima, Laura C. Brown, Dor Salomon, Julia C. van Kessel

**Author notes:** Corresponding author: Julia van Kessel. Emra Klempic: Loretto Health and Rehabilitation, Syracuse, NY, 13205.

## Abstract

The bacterial pathogen *Vibrio coralliilyticus* (*Vcor*) causes disease in coral species worldwide. The mechanisms of *Vcor* coral colonization, coral microbiome interactions, and virulence factor production are understudied. In other model *Vibrio* species, virulence factors like biofilm formation, toxin secretion, and protease production are controlled through a density-dependent communication system called quorum sensing (QS). Comparative genomics indicated that *V. coralliilyticus* genomes share high sequence identity for most of the QS signaling and regulatory components identified in other *Vibrio* species. Here, we identify an active QS signaling pathway in two *V. coralliilyticus* strains with distinct infection etiologies: type strain BAA-450 and coral isolate OCN008. The inter-species AI-2 autoinducer signaling pathway in both strains controls expression of the master QS transcription factor VcpR to regulate >300 genes, including protease production, biofilm formation, and two conserved type VI secretion systems (T6SSs). Activation of T6SS1 by QS results in secretion of effectors and enables interbacterial competition and killing of prey bacteria. We conclude that the QS system in *V. coralliilyticus* is functional and controls expression of genes involved in relevant bacterial behaviors that may influence coral infection.

**IMPORTANCE:** *Vibrio coralliilyticus* infects many marine organisms, including multiple species of corals, and is a primary causative agent of tissue loss diseases and bacterial-induced bleaching. Here we investigate a common cell-cell communication mechanism called quorum sensing, which is known to be intimately connected to virulence in other *Vibrio* species. Our genetic and chemical studies of *V. coralliilyticus* quorum sensing uncovered an active pathway that directly regulates key virulence factors: proteases, biofilms, and secretion systems. These findings connect bacterial signaling in communities to infection of corals, which may lead to novel treatments and earlier diagnoses of coral diseases in reefs.

## INTRODUCTION

*Vibrio coralliilyticus* (*Vcor*) is a Gram-negative bacterium and prolific marine pathogen. Since the description of this species in the early 2000s (1), *Vcor* has remained an important etiological agent in aquacultural industries and along global coastlines. This is, in part, due to the range of hosts affected by this species, which includes bivalve larvae (2–5), fish (6), urchins (7), and various coral species (8–13). Acute environmental changes can result in the disruption of the complex processes and symbioses formed among a coral’s microbiota community, which can results in disease under certain conditions (14, 15). The coral microbiota is a diverse community encompassing fungi, bacteria, microeukaryotes, archaea, viruses, and for many coral species the endosymbiont family, *Symbiodiniaceae* (16, 17). When corals are infected by *Vcor*, it can result in bleaching (loss of the endosymbiotic dinoflagellates) or tissue loss (destruction of the healthy tissue) depending on the pathogenic strain and environmental conditions (10, 18–22). While disease itself is natural for any system, disease outbreaks, when disease prevalence increases above the normal baselines, can result in mass coral mortalities that have larger ramifications for local biodiversity and the economy of surrounding communities (23–26). Unfortunately, the specific mechanisms behind the initiation of coral disease outbreaks or the factors that sustain an outbreak are unknown. While elevated water temperatures are important for virulence of some *Vcor* strains (10, 21, 22, 27), other environmental factors are unclear. The genetic regulation and identity of virulence factors responsible for many systems in *Vcor* have yet to be fully explored. Current knowledge links one regulatory pathway of *Vcor*’s pathogenicity to the bacterial phenomenon known as quorum sensing (QS) (18, 28).

QS is a cell-to-cell communication system utilized by bacteria to control behaviors in a density-dependent manner (29–33). In *Vibrio* species, these behaviors broadly include competence, swarming motility, bioluminescence, biofilm formation, type III secretion system (T3SS), and type VI secretion system (T6SS) activity (30, 31, 34–37). The QS signal-transduction circuit and regulatory network has been characterized in several *Vibrio* species (38, 39), including *Vibrio cholerae* (40, 41)*, Vibrio campbellii* (42–44)*, Vibrio parahaemolyticus* (45), *Vibrio alginolyticus* (46), *Vibrio fischeri* (47), and others. This collective body of research yields a strong foundation to test and compare QS in newly identified *Vibrio* strains. *Vibrio* QS systems rely on signal transduction networks comprising small molecule autoinducer (AI) synthases, cognate hybrid histidine kinase (HK) receptor proteins, a phosphotransfer protein, a response regulator, small regulatory RNAs (sRNAs), and transcriptional regulators. The broadly defined QS system studied in the *Vibrio* genus is depicted in Figure 1, with the predicted *Vcor* gene names indicated. The known AI synthases of the *Vibrio* genus include LuxM, CqsA, and LuxS that produce autoinducer-1 (AI-1), cholera autoinducer-1 (CAI-1), and autoinducer-2 (AI-2), respectively, which diffuse out of the cell and into the external environment. At low cell density (LCD), AIs are present at insufficient concentrations in the extracellular environment to bind to their respective cognate HK membrane-bound receptor proteins, LuxN, CqsS, and LuxPQ, respectively. There are also two HK receptors recently identified in *V. cholerae*, CqsR and VpsS, with unknown cognate AI signals that function in this pathway (48, 49). Further, the cytoplasmic HqsK/H-NOX receptor binds nitric oxide (NO) and is yet another functional HK in the pathway (49, 50). Each of these receptor proteins (LuxN, CqsS, LuxPQ, CqsR, HqsK) are either predicted or have been shown to function as kinases in the absence of cognate ligands and autophosphorylate at their conserved histidine residue, transfer the phosphate group to the aspartic acid in the receiver domain, then transfer the phosphate to the histidine of the LuxU phosphotransfer protein (present in all *Vibrio* sp.), and finally to the aspartic acid of the LuxO response regulator (Fig. 1). Phosphorylated LuxO, together with Sigma-54, activates the transcription of the quorum regulatory (Qrr) sRNAs, which together with Hfq degrade the transcript of the TetR-type master transcriptional regulator, for example LuxR in *V. campbellii*, but activate the translation of AphA to further transcriptionally regulate LCD bacterial behaviors (Fig. 1). At high cell density (HCD), extracellular AI concentrations are high, leading to binding of AIs to their cognate receptors at the membrane or in the cytoplasm. AI-binding switches these HKs from functioning as kinases to phosphatases, thus reversing the flow of phosphate upstream of LuxO. As a result, LuxR expression levels increase, AphA production ceases, and LuxR functions as a master transcriptional regulator activating the expression of HCD group behaviors (Fig. 1). A separate QS signaling system identified in *V. cholerae* involves the synthase protein, Tdh, that produces the molecule DPO, which interacts with cytoplasmic VqmA to activate the transcription of VqmR sRNA to inhibit biofilm formation.

**Figure 1.**
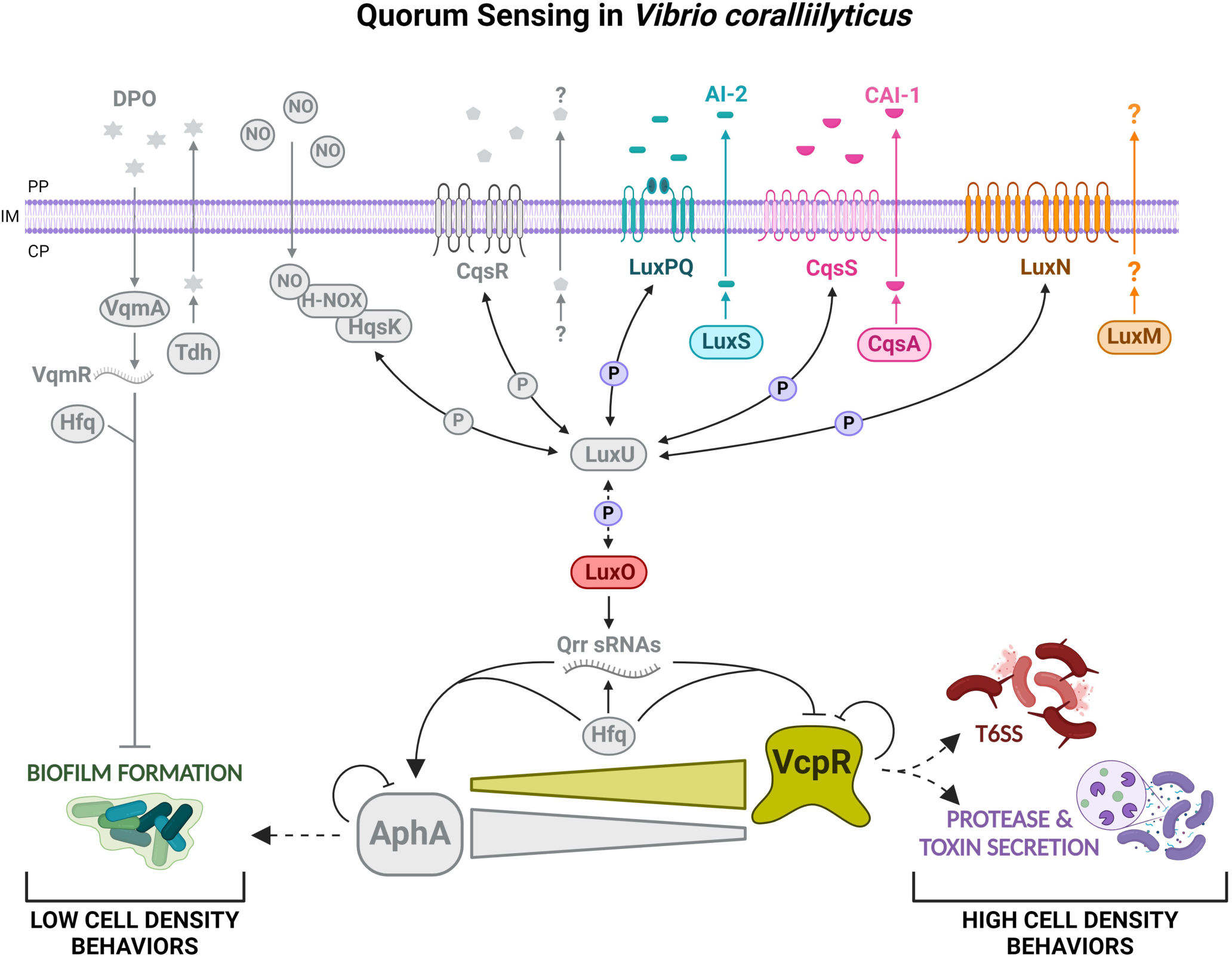
Model of the quorum sensing regulatory pathway in *Vcor*. The putative QS system of *Vcor* and proteins predicted to function in the QS circuit are shown. Bright colored proteins represent experimentally confirmed pathways either in this manuscript or previously published research. Gray colored proteins and pathways have yet to be investigated in *Vcor*. The predicted histidine kinase binding proteins (LuxPQ, CqsS, LuxN, CqsR, and HqsK) encoded by *Vcor* function as phosphatases at high cell density and kinases at low cell density; these proteins transfer phosphate to and from two core QS regulatory proteins LuxU and the final phosphate relay protein, LuxO. At HCD, dephosphorylated LuxO inactivates the transcription of the Qrr sRNAs, and the protein levels of the VcpR master transcriptional regulator increase to regulate its own feedback loop and group behaviors (e.g., type VI secretion system [T6SS], protease and toxin secretion). At LCD, transcribed Qrr sRNAs block production of VcpR and activate the translation of the transcriptional regulator AphA, leading to the regulation of individual bacterial behaviors (e.g., biofilm formation). Image created on BioRender.com.

To provide context and delineate between quorum sensing systems in the field, the LuxR protein described in *V. campbellii* (previously classified as *Vibrio harveyi* (51)) has no functional or genetic similarity to the LuxR family of proteins that bind and respond to acyl-homoserine lactone (AHL) AIs produced by LuxI in the *V. fischeri* system. LuxR from *V. campbellii* was named due to its regulation of *luxCDABE*, but it was noted by the researchers at the time of discovery that these two types of proteins are not the same: “The amino acid sequence of the LuxR product of *V. campbellii*, which indicates a structural relationship to some DNA-binding proteins, is not similar to the sequence of the protein that regulates expression of luminescence in *V. fischeri*.”, stated in the abstract of the study by Showalter *et al.* (30, 52).

Although *Vcor* was first described only ∼20 years ago, the field has uncovered several important virulence factors and regulatory networks that play a role in coral disease. Research completed in *V. tubiashii* type strain RE22, later reclassified as *Vcor* (2, 53), identified VcpR (then called VtpR) as a TetR-type protein and global transcriptional QS regulator (54). VcpR shares ∼84% amino acid identity with *V. campbellii’s* LuxR (54). Subsequent studies uncovered that VcpR regulates the expression of hemolysins and at least two metalloprotease genes, *vcpA* and *vcpB*, suspected to be critical virulence factors for bivalve larvae (54, 55). Three *Vcor* strains have been studied to determine molecular mechanisms of infection: type strain BAA-450 (=ATCC BAA-450 (56), =YB1 (8), =LMG 20984 (6)), OCN008, and OCN014, which have all been demonstrated to infect corals and larval oysters in both temperature- and dose-dependent manners (2, 9, 27). These responses to increased temperatures or conditions that promote *V*cor growth suggest a link between specific environmental conditions contributing to disease outbreaks (57, 58). For example, increased seawater surface temperature (SST) triggers virulence in *Vcor*. Depending on the host, different virulence factors are upregulated at higher water temperatures above 27°C in strains BAA-450 and OCN014, whereas OCN008 virulence is active at lower and higher temperatures (from 23°C to above 27°C) (1, 10, 27, 56, 59, 60).

Each of these strains exhibits cross-species infection by utilizing a core set of cell-associated virulence factors, the transcriptional regulator ToxR, and putative adherence factor OmpU (27). It is hypothesized strains OCN008, OCN014, and BAA-450 have strain-specific etiologies because they may have evolved in different environments and in different hosts (1, 8–10, 61, 62).

To characterize the QS signaling system in *Vcor*, our group sequenced the genome of *Vcor* OCN008 (63) and performed a comparative genomics analysis to determine the putative QS system components conserved in *Vcor* strains OCN008, BAA-450, and OCN014. We investigated the presence of AIs, receptors, and transcription factors. Collectively, we show that the AI-2 synthesis and sensing system is intact and active, though there are likely other signal(s) active as well. We determined the VcpR regulons for two strains, BAA-450 and OCN008, and determined that both strains activate multiple virulence factors, including a T6SS and proteases, at high cell densities. From this study, we conclude that QS functions to integrate environmental signals of cell density and species identity to control expression of virulence genes that impact coral infection.

## RESULTS

### Vcor encodes homologs of quorum sensing proteins

In *Vcor,* VcpR is a known functional homolog of the LuxR/HapR-type master transcriptional regulators characterized in other *Vibrio* QS systems (64). Because VcpR has been shown to regulate gene expression in *Vcor*, we hypothesized that *Vcor* encodes a functional QS system. We used comparative genomics and examined fourteen fully sequenced *Vcor* genomes in Genbank. Using a previously assembled phylogenetic tree of the *Vibrio* clade (42), we examined the conservation of genes encoding autoinducer synthases, autoinducer receptors, response regulators, and transcription factors in the QS circuit. The query sequences were chosen from the model QS strain *V. campbellii* BB120 (Fig. S1). Using this method, any proteins with amino acid identity below 40% were not designated as a homolog and appear as black on the heat-map (Fig. S1). After performing this initial analysis to identify putative QS system genes in *Vcor*, we used the *Vcor* OCN008 strain genes as query sequences to examine conservation among *Vcor* genomes (Fig. 2A). As expected based on published experimental data in the *Vcor* field (64–66), VcpR was highly conserved among most *Vcor* strains, sharing 85% amino acid identity with BB120 LuxR (Fig. 2A). However, VcpR appeared to be poorly conserved in strain SNUTY-1, sharing about 67% amino acid identity with strain OCN008 (Fig. 2A). Other highly conserved QS genes included LuxO, LuxU, Hfq, and AphA (Fig. 2A). Most of these *Vcor* strains also encoded homologs of at least one pair of autoinducer receptors and synthases. In addition, bioinformatics confirmed the conservation of synthase protein, Tdh, and cytoplasmic regulatory protein, VqmA. Notably, the autoinducer-1 (AI-1) synthase LuxM is poorly conserved in many of the *Vcor* genomes (Fig. 2A). However, for several strains, the presence of a gene product with a higher amino acid identity with BB120 LuxM correlated with the presence of a LuxN homolog with increased amino acid identity to *V. campbellii* BB120 (Fig. 2A). Surprisingly, in some strains such as OCN008 and BAA-450, there were two LuxN paralogs proximally encoded at one genetic locus, though these were only ∼46% identical to each other (Fig. 2B; Table S4). None of these 14 *Vcor* strains encoded a VpsS homolog sharing amino acid identity above 40% with BB120 VpsS (Fig. S1). From these data, we conclude that most of these 14 sequenced *Vcor* strains encode a suite of proteins that could constitute a complete QS signaling circuit with the potential to produce, sense, and respond to at least one autoinducer.

**Figure 2.**
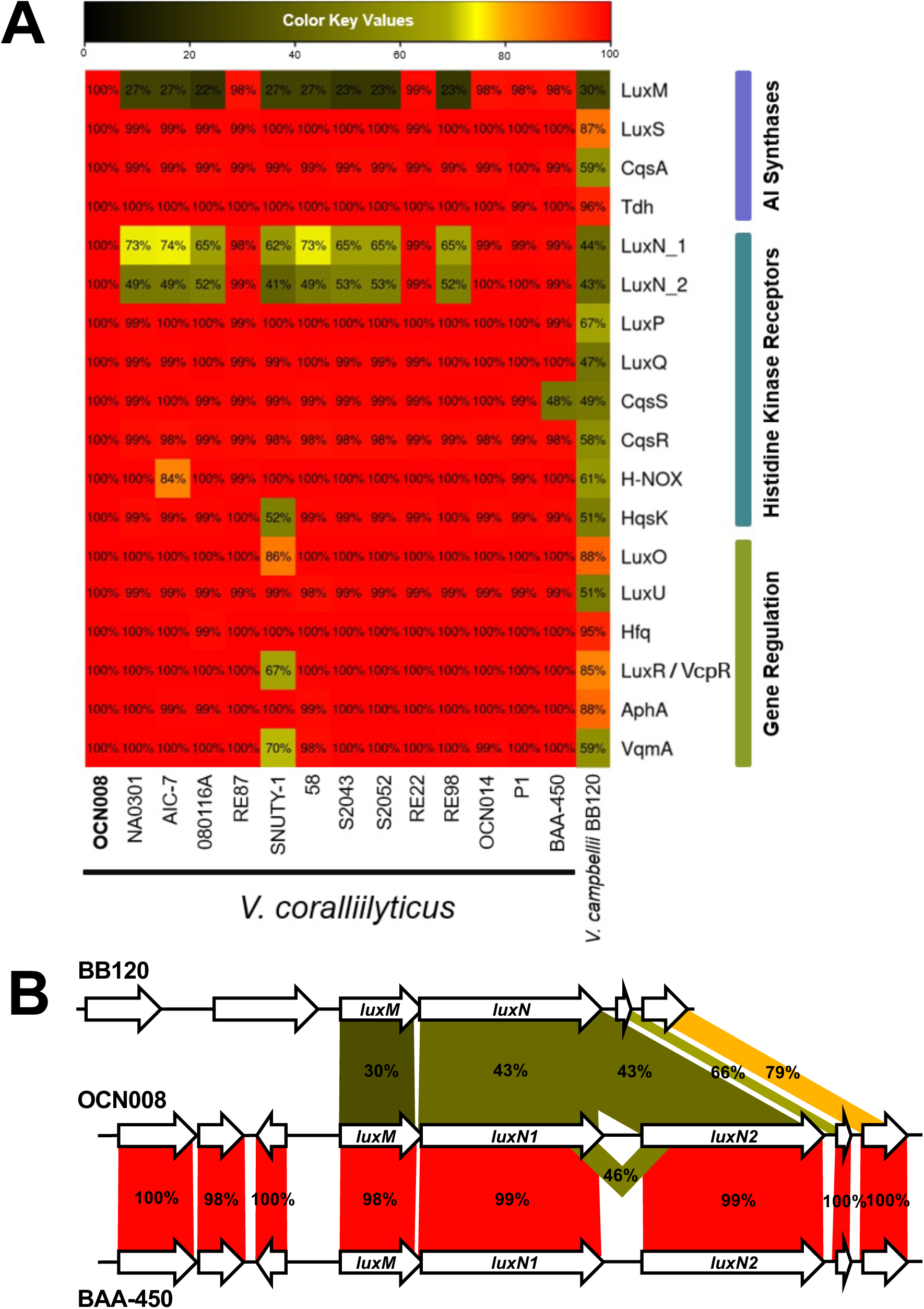
Conservation of QS system homologs in *Vcor* strains. (A) The proteins listed on the y-axis are involved in QS in *V. campbellii* BB120 and are grouped based on function. The heat-map indicates amino acid conservation of these proteins in *Vcor* strains using OCN008 genes as query sequences. (B) Genetic loci in *V. campbellii* BB120 and *Vcor* OCN008 and BAA-450 strains. Percent amino acid identity is indicated for the pair-wise comparisons.

### VcpR expression responds to changes in cell density and QS signaling

We previously established a reliable reporter for transcriptional regulation by the family of LuxR-type proteins that utilizes the *luxCDABE* operon (43, 48, 49, 67, 68). Although this is a heterologous reporter, its use in numerous *Vibrio* species has been verified as a *bona fide* reporter for LuxR activity in *Vibrio* species (48, 69). These previous studies showed that binding of a LuxR-type protein strongly and specifically activates this promoter with a dynamic range of more than three orders of magnitude. Because LuxR production is determined by Qrr activation, expression of *luxCDABE* and bioluminescence production correlate with production and detection of autoinducers in a culture. To test the transcriptional function of VcpR using this reporter, we introduced the *luxCDABE* reporter plasmid (pCS18) into wild-type strains and Δ*vcpR* mutant strains of OCN008, OCN014, and BAA-450. We measured bioluminescence (Lux/OD_600_) production over time (OD_600_) for *Vcor* strains containing pCS18. We observed the predicted U-shaped curve that is typical for *Vibrio* strains expressing bioluminescence in a QS-dependent manner: cells at HCD (from the inoculum) produced high levels of bioluminescence, which was diluted out by cell growth in the initial hours of a growth curve. Cells produced very little light at LCD because there were low levels of autoinducers. As the cells grew and reached quorum (OD_600_ = ∼0.3), light production sharply increased and accumulated to maximum levels at HCD at OD_600_ = ∼1.0 in all three *Vcor* strains containing pCS18 (Fig. 3A). Comparatively, the Δ*vcpR* strains containing pCS18 produced no bioluminescence and were constitutively “dark” throughout the curve, similar to wild-type strains that do not contain the pCS18 reporter (Fig. 3A).

**Figure 3.**
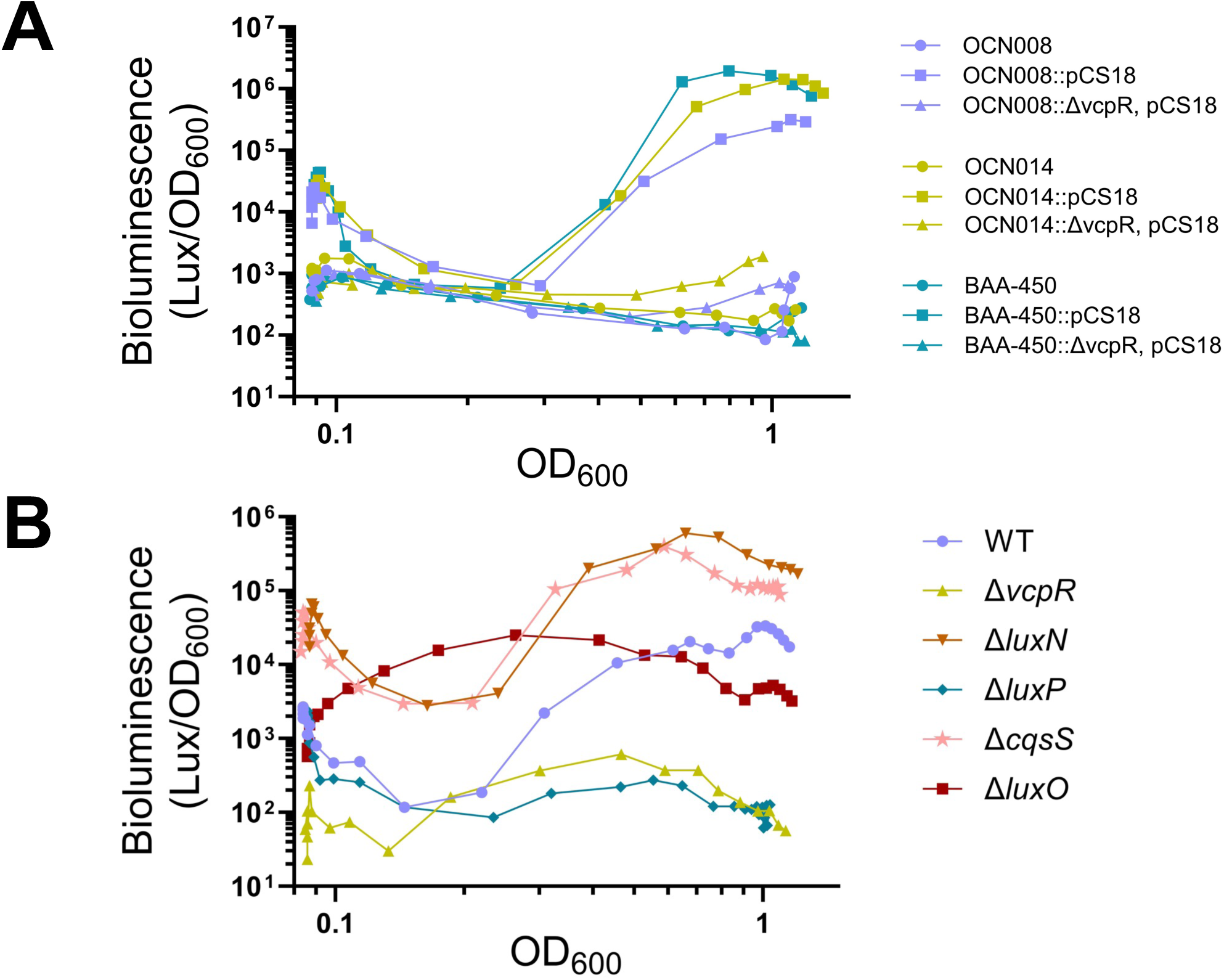
*Vcor* responds to changes in cell density. (A) Bioluminescence production in *Vcor* OCN008, OCN014, and BAA-450 wild-type and Δ*vcpR* reporter strains (wild-type, wild-type::pCS18, and Δ*vcpR*::pCS18). (B) Bioluminescence production in OCN008 reporter strains (wild-type and mutant strains). For panel A, cultures were grown in LB marine broth and for panel B, cultures were grown in minimal media (M9) salts supplemented with glucose and casaminoacids. Bioluminescence data was normalized to cell density (OD_600_) over the course of the growth curve. The data shown are from a single experiment and are representative of three biological replicate assays.

To test the function of putative QS circuit homologs in *Vcor*, we focused on strain OCN008 and deleted a subset of the putative *Vcor*-encoded QS genes identified by bioinformatics and assayed their function using bioluminescence. As observed in other *Vibrio* species, deletion of *luxO* resulted in a constitutive bioluminescent phenotype (Fig. 3B) (42, 48, 49, 70, 71). Deletion of the genes encoding two putative AI receptor histidine kinases, CqsS and LuxN2, resulted in increased bioluminescence throughout the curve, indicating that both proteins act to repress QS signaling at LCD (Fig. 3B). We also observed that a Δ*luxP* deletion strain exhibited low bioluminescence throughout the curve (Fig. 3B). In *V. campbellii*, LuxP is a periplasmic binding protein that controls the quaternary state of LuxQ and its kinase activity based on binding of AI-2 (72). In the Δ*luxP* strain, we observed a phenotype mimicking a locked LCD-phenotype, suggesting that LuxQ was constitutively kinase-active (Fig. 3B). Conversely, deletion of *luxN* or *cqsS* resulted in increased overall bioluminescence throughout the curve (49, 73). However, even with increased bioluminescence compared to wild-type, both the Δ*luxN* and Δ*cqsS* strains responded to density, indicating other inputs into the system exist. From these data, we conclude that *Vcor* encodes QS proteins LuxO, CqsS, LuxN, and LuxP that display analogous functions to those defined in other *Vibrio* species with regard to QS signal response and regulation.

### The Vcor QS circuit regulates biofilm formation and protease production

We next investigated the role of QS signaling in regulation of other physiological processes that are typically controlled by QS in *Vibrio* species: biofilm and protease production (41, 45, 46, 74). Biofilm production has previously been shown to be a LCD behavior in *V. cholerae* and is repressed by HapR, another LuxR family protein (75, 76). At LCD, nonmotile bacteria excrete extracellular *Vibrio* polysaccharides (VPS), which surround the cell and form an extracellular matrix (75). As the cells grow toward HCD, *V. cholerae* HapR represses expression of VPS and its activators (77). In addition, AphA positively regulates biofilm formation at LCD (78, 79). This pattern is similar in other *Vibrio* species (80, 81). If this model holds true for *Vcor* QS, we predicted that biofilm production would increase in strains with mutant alleles that increase LuxO phosphorylation and Qrr transcription (*e.g.*, Δ*luxP*), resulting in lower levels of VcpR and higher levels of AphA. Similarly, we predicted that biofilm formation would be lowest in strains unable to activate the Qrrs (*e.g.*, Δ*luxO*), resulting in high levels of VcpR and low levels of AphA.

Using our set of gene deletion strains in OCN008, we assayed biofilm formation using a standard crystal violet assay. Wild-type OCN008 produced low levels of biofilms, which increased ∼2-fold in a Δ*vcpR* mutant but was not significantly different than wild-type (Fig. 4A). Similarly to the wild-type phenotype, the HCD-locked mutant, Δ*luxO*, resulted in low levels of biofilm production (Fig. 4A). We observed that the Δ*luxP* mutant had the largest biofilm production (Fig. 4A), which aligned with our previous prediction that this strain has constitutive LuxQ kinase activity that drives a LCD-locked phenotype.

**Figure 4.**
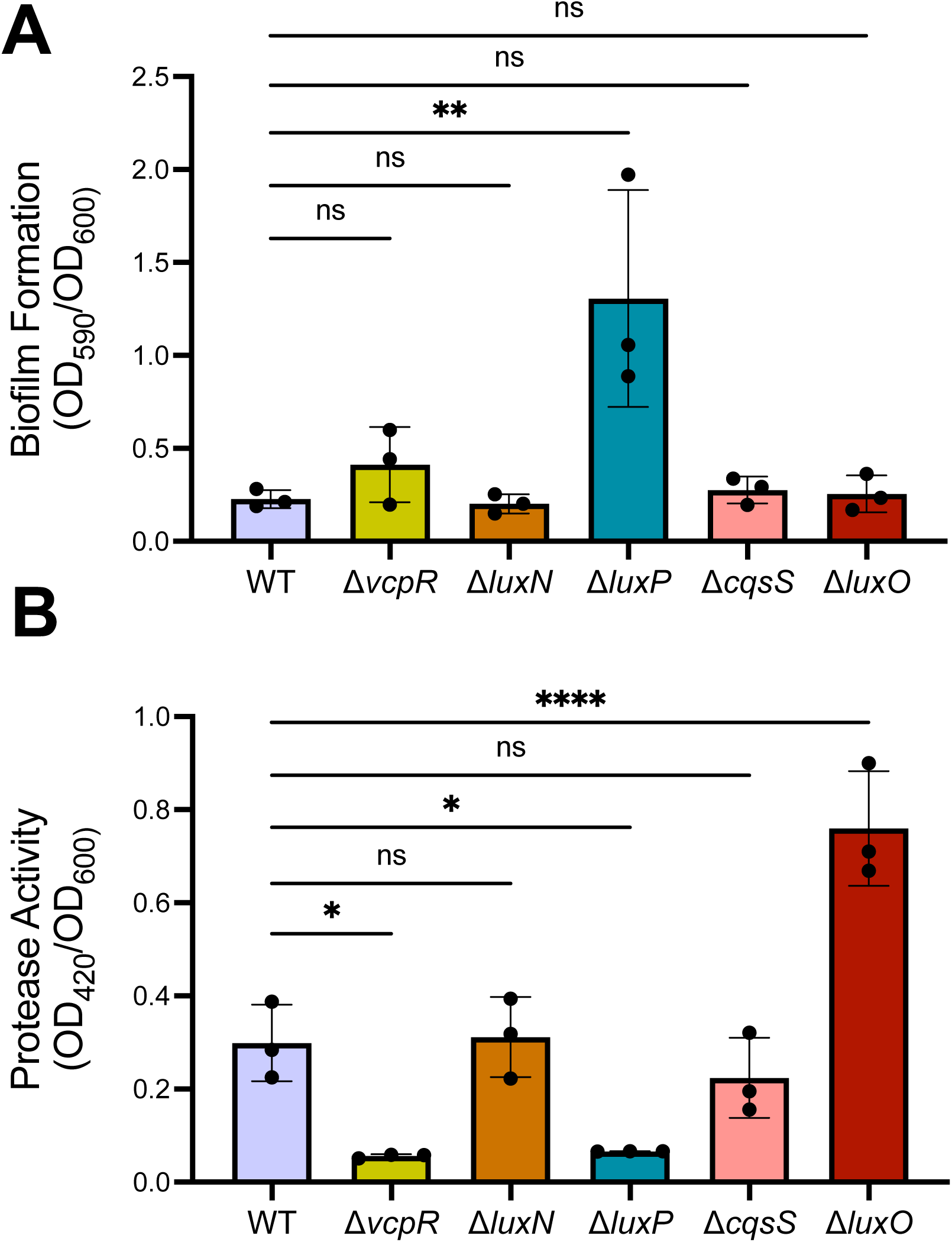
Biofilm production and protease activity in *Vcor* wild-type and mutant strains. (A) Biofilm production measured using crystal violet in *Vcor* wild-type strain OCN008 and gene deletion strains. (B) Protease activity in *Vcor* wild-type strain OCN008 and gene deletion strains. Asterisks indicate significance (one-way analysis of variance; * = *p*<0.05, ** = *p*<0.01, **** = *p*<0.0001, ns = not significant; *n* = 3 for each assay) on normally distributed data (Shapiro-Wilk test) followed by Dunnett’s multiple comparisons test.

In several other *Vibrio* species, proteases are produced at HCD and have important roles in host infection (65, 82, 83). Similarly, *Vcor* produces proteases, some of which are known to be regulated by VcpR. To examine the effects of QS signaling on protease production, we assayed knockout strains in a protease assay that measures digestion of azocasein via a colorimetric assay. We observed the lowest protease activity in both the Δ*vcpR* and Δ*luxP* mutant strains. Wild-type OCN008, Δ*luxN,* and Δ*cqsS* strains all displayed similar levels of protease activity (Fig. 4B), whereas the Δ*luxO* strain produced significantly more protease activity than wild-type. The absence of biofilm and protease phenotypes distinguishable from wild-type for the Δ*luxN* and Δ*cqsS* strains was inconclusive; in other *Vibrio* species, deletion of a single kinase does not always have a phenotype due to the activity of other kinases in the system (49). From these results, we conclude that biofilm formation is repressed by QS at HCD, and protease activity is activated by QS at HCD. In addition, these two behaviors are regulated at least by VcpR and LuxP.

### V. coralliilyticus produces AI-2

The majority of sequenced *Vcor* genomes appeared to encode homologs of AI synthases LuxS and CqsA (Fig. 2A) (42). Some strains, such as *Vcor* OCN008 and BAA-450, do encode LuxM homologs but these predicted proteins are very different than LuxM in BB120 (Fig. 2B). Thus, we predicted that if AIs are produced by these LuxM proteins, the structures may be different than HAI-1 produced by BB120 LuxM. We also predicted that *Vcor* strains likely produced AI-2 and CAI-1 due to the high level of conservation of LuxS and CqsA, respectively.

To investigate the AIs produced by *Vcor*, we used a combination of mass spectrometry and *in vivo* reporter strain analyses. First, we examined production of AHLs by *Vcor* strains OCN008, OCN014, and BAA-450*. V. campbellii* BB120 served as a positive control because it produces the HAI-1 molecule, which is readily identifiable by mass spectrometry, as are numerous synthetic AHL compounds (42). However, using the same methods we have used previously to identify AHLs, we were unable to detect any AHLs in the supernatant extracts of the three *Vcor* strains (Fig. S2). In addition, we used an *in vivo* reporter assay in which the *V. campbellii* strain BB120 produced bioluminescence in response to supernatants containing AIs. Isogenic BB120 strains containing deletions of AI synthases thus serve as reporters for the presence of AI in the supernatant of test samples. In this experiment, we added supernatants of BB120, OCN008, OCN014, and BAA-450 to a Δ*luxM* strain of *V. campbellii* and observed that only the BB120 supernatant produced bioluminescence above the media-only control (Fig. 5A). From these negative data, we cannot conclude whether the three *Vcor* strains tested produce AHLs; the levels of AI might be too low to detect, or the AHL produced may have compositional differences that prohibit detection by mass spectrometry or our *in vivo* reporter.

**Figure 5.**
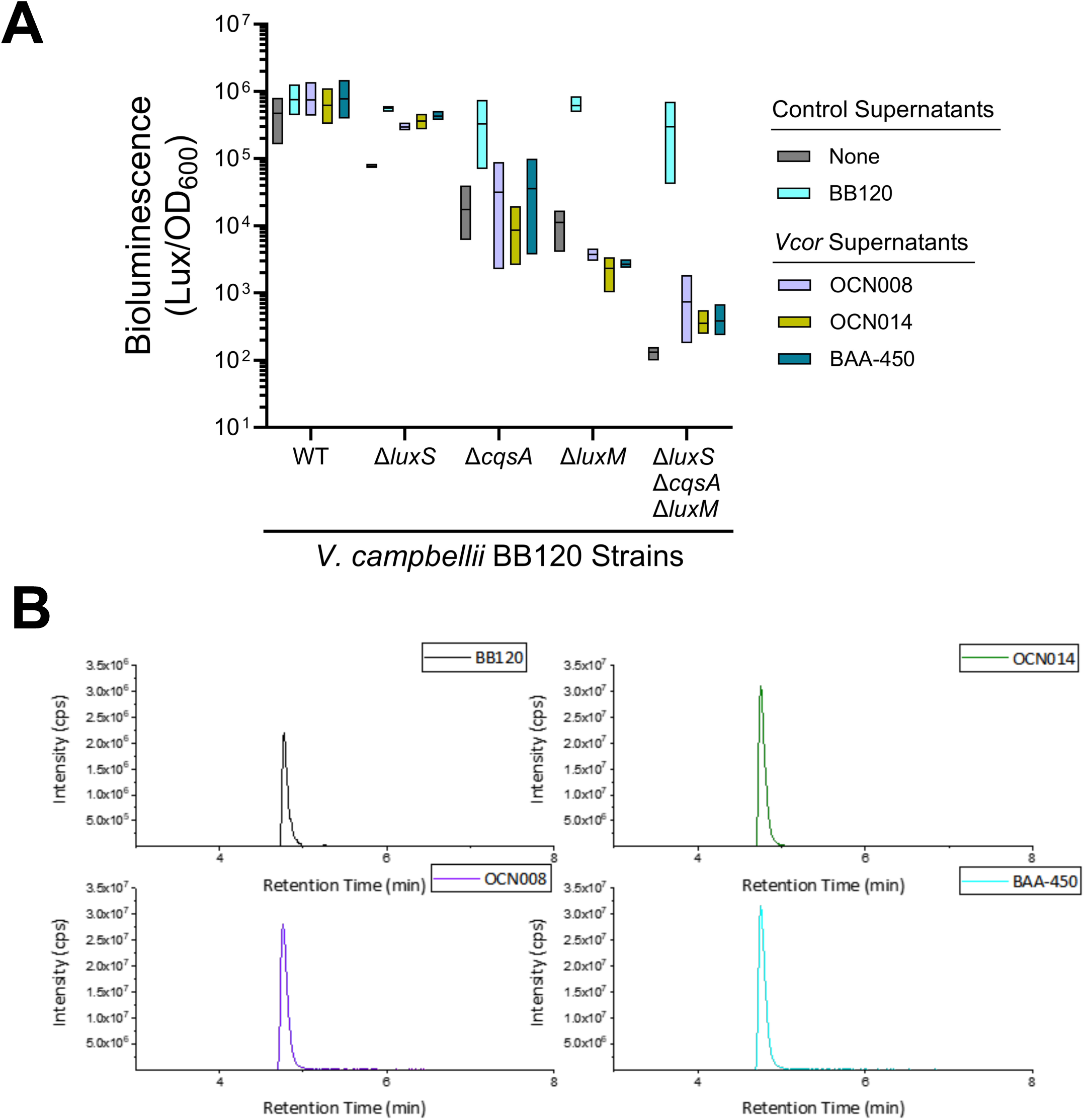
*Vcor* produces AI-2. (A) Bioluminescence production normalized to cell density (OD_600_) in *V. campbellii* BB120 type strains after g supernatant from *V. campbellii* BB120 and *Vcor* OCN008, BAA-450, and OCN014 (n = 3). (B) Extracted ion chromatograms from supernatant extracts of *V. campbellii* BB120 (positive control) and *Vcor* OCN008, BAA-450, and OCN014 depict the peak corresponding to the positive detection of AI-2.

Next, we examined production of AI-2 by *Vcor* strains. In other *Vibrio* strains, AI-2 is synthesized by LuxS as a metabolic byproduct called DPD, or 4,5-Dihydroxy-2,3-pentanedione (84). DPD reacts enzymatically with boric acid to form a furanosyl borate diester (Fig. S3A), which is the compound specifically recognized by the membrane receptor LuxPQ (85). We utilized a chemical derivatization to measure DPD levels in supernatants, which formed the compound DPDQ (Fig. S3B). Mass spectrometry readily identified DPDQ in the supernatants of all three *Vcor* strains at similar levels to that produced by BB120 (Fig. 5B). Further, *Vcor* supernatants added to the Δ*luxS* BB120 reporter resulted in bioluminescence production similar to that of BB120 supernatant (Fig. 5A). From these data, we conclude that *Vcor* strains OCN008, OCN014, and BAA-450 produce AI-2 that can be detected by BB120.

Third, we examined production of the fatty acid-like AI synthesized by CqsA by *Vcor* strains. *V. cholerae* CqsA produces CAI-1 ((*S*)-3-hydroxytridecan-4-one), whereas *V. campbellii* BB120 produces an enamine variant of CAI-1 [*(Z)*-3-aminoundec-2-en-4-one (Ea-C8-CAI-1)] (86). We were able to detect synthetic CAI-1 using mass spectrometry, but we did not detect CAI-1 in *V. cholerae* supernatants (data not shown). We suspect this may be a limitation of the quantity of supernatants we extracted. However, we were able to detect the enamine-CAI-1 in the BB120 supernatant (Fig. S2B). We did not detect either compound from the three *Vcor* supernatants tested. Further, the *in vivo* reporter assay indicated that the *Vcor* supernatants did not contain sufficient enamine-CAI-1 to result in bioluminescence expression in the Δ*cqsA* BB120 strain (Fig. 5B). From these negative data, we cannot conclude whether *Vcor* produces a CAI-1-like molecule, for the same reasons given above for AHL.

### VcpR regulons in BAA-450 and OCN008

We next determined the VcpR regulons in strains BAA-450 and OCN008 using RNA-seq comparing wild-type and Δ*vcpR* isogenic strains in OCN008 and BAA-450. In OCN008, VcpR regulated 896 genes 2-fold or more (false-discovery rate 0.05), whereas in BAA-450 the VcpR regulon was 363 genes (Tables S5, S6). Both strains showed significant VcpR-dependent activation of genes in the T6SS1 (BAA-450: VIC_RS16230-360) and T6SS2 (BAA-450: VIC_RS26410-150), two distinct protein secretion systems encoded in both strains (see section below), which were recently implicated in virulence (87), with a larger difference in VcpR-dependent expression for T6SS1 genes (Fig. 6). Additional genes were detected as significantly positively regulated by VcpR that are correlated with virulence. For example, the protease-encoding genes *vcpA* and *vcpB* were both activated by VcpR, both of which we verified using RT-qPCR (Fig. S4). In addition, *vpsR* was repressed by VcpR in OCN008 but not in BAA-450 (Tables S5, S6). In *V. cholerae*, the master biofilm regulator VpsR binds cyclic di-GMP and activates transcription of the biofilm operons (88). OCN008 VcpR also repressed VpsT, though less than 2-fold. These results are consistent with the slight but not significant increase in biofilm formation in the Δ*vcpR* strain compared to wild-type (Fig. 4A). These findings also align with the literature showing that multiple regulators control biofilm formation in addition to VcpR at LCD (*e.g.*, sRNAs, AphA), which likely explains why the Δ*luxP* mutant biofilm phenotype is significantly different than wild-type because LuxP is upstream of VcpR, AphA, and sRNAs in the pathway (88). From these results, we conclude that both *Vcor* strains OCN008 and BAA-450 broadly regulate genes via VcpR, though to different extents.

**Figure 6.**
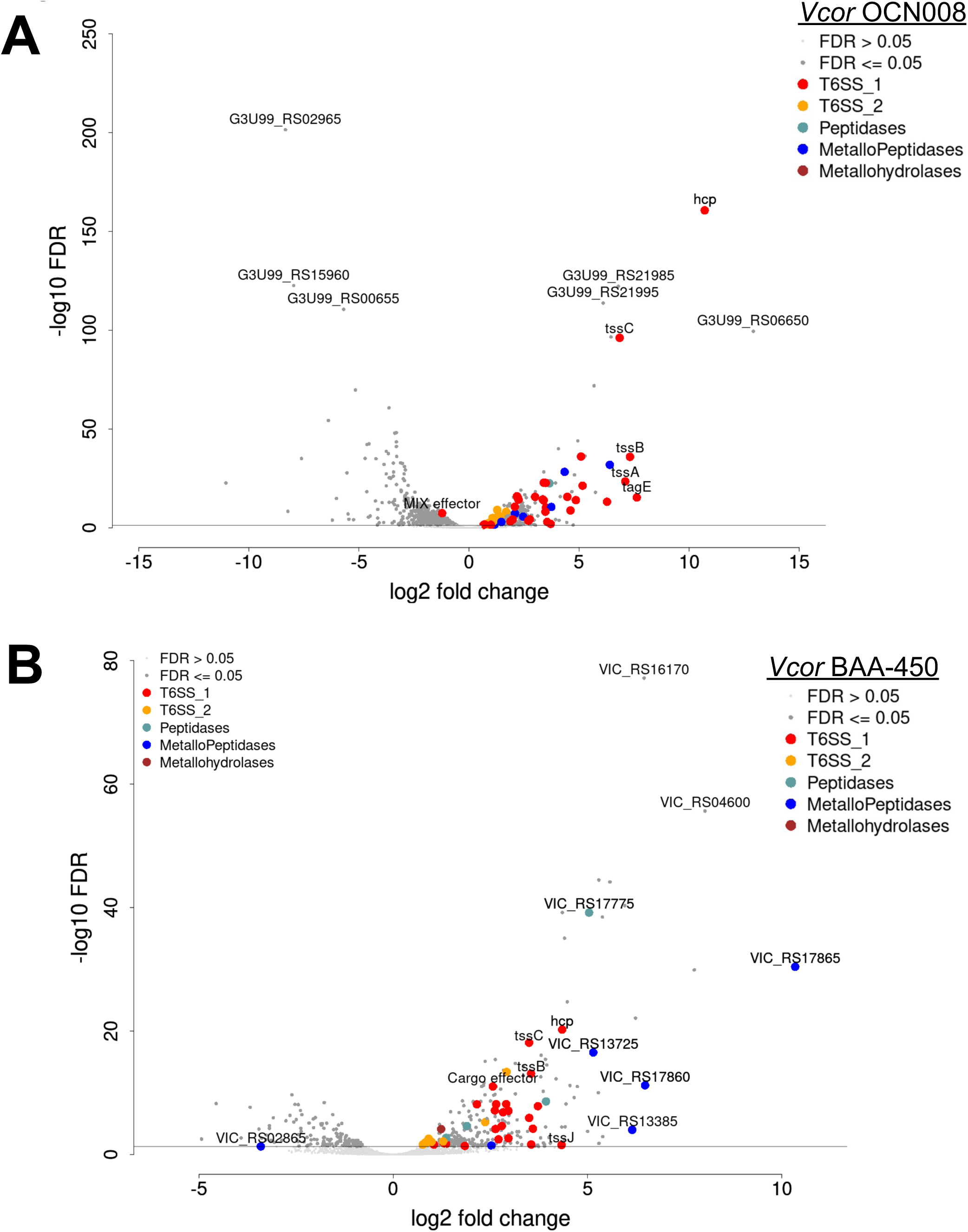
VcpR regulates virulence factors predicted to be involved in pathogenesis. Volcano plots of the RNA-seq data are shown for (A) OCN008 wild-type versus Δ*vcpR* and (B) BAA-450 wild-type versus Δ*vcpR*.

### QS activates T6SS1 secretion and represses T6SS2 secretion

T6SS1 has been shown to be active in *Vcor* OCN008, and VcpR was shown to be necessary for activation of the system and for T6SS1-mediated interbacterial competition (36). Recently, we showed that there are two active T6SSs encoded by *Vcor* strains: T6SS1, which secretes mostly antibacterial effectors and plays a role in interbacterial competition, and T6SS2, which secretes an arsenal of novel anti-eukaryotic effectors and directly contributes to virulence Additionally, our RNA-seq analysis (see section above) and a *Vcor* secretome analysis identified genes outside of the main T6SS1 and T6SS2 gene clusters that encode structural T6SS components, effectors, and immunity proteins (87) (Table S7). To investigate the role of QS signaling in the activation of both systems in strain OCN008, we monitored the expression and secretion of the T6SS1 and T6SS2 core components VgrG1 and Hcp2, respectively. The Δ*hcp1* and Δ*tssM2* strains served as negative controls for T6SS1 and T6SS2 activity, respectively (87). We observed that the wild-type strain expressed and secreted VgrG1, whereas mutant strains that mimic LCD-phenotypes (i.e., Δ*luxP* and Δ*vcpR*) did not express detectable levels of VgrG1 (Fig. 7), suggesting that T6SS1 is inactive at low cell density. This result aligns with the observed ∼10-fold decrease in *vgrG1* gene expression via RNA-seq in the OCN008 Δ*vcpR* strain compared to wild-type (Table S5). As expected, the Δ*hcp1* strain expressed VgrG1 but did not secrete it.

**Figure 7.**
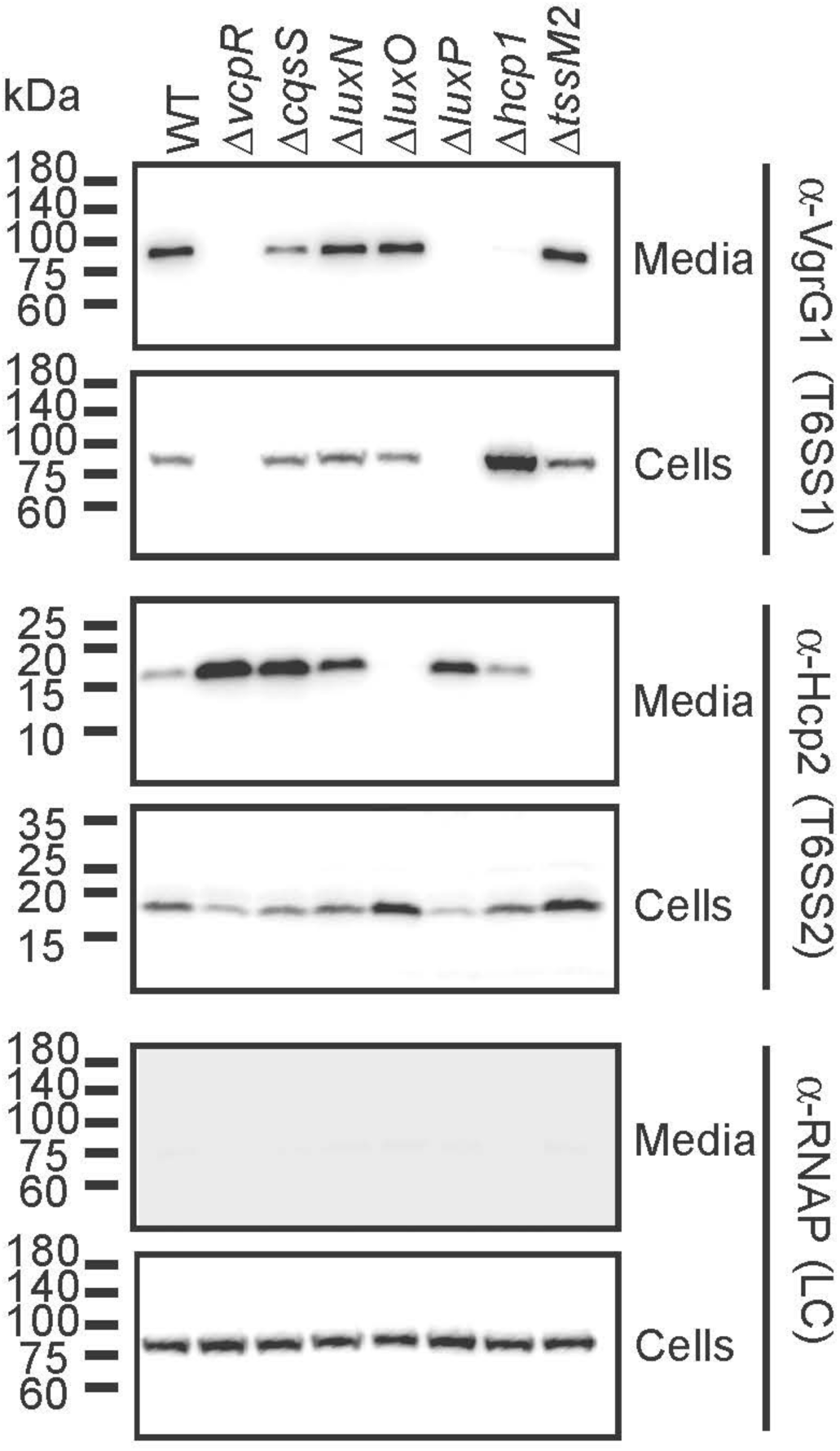
T6SS1 and T6SS2 are differentially regulated by QS through VcpR. Expression (cells) and secretion (media) of VgrG1 and Hcp2, indicating T6SS1 and T6SS2 activity, respectively, from the indicated *Vcor* OCN008 strains. RNA polymerase sigma 70 (RNAp) was used as a loading and lysis control. Results from a representative experiment out of three independent experiments are shown.

In contrast to T6SS1, secretion via T6SS2 was mildly induced in the mutant strains that mimic LCD (Fig. 7). This result contrasts with the RNA-seq data in which *hcp2* expression was decreased ∼3-fold in the Δ*vcpR* strain (Table S5). In addition, the HCD-locked strain Δ*luxO* was defective in secretion of Hcp2 from T6SS2 but not in production of the Hcp2 protein (Fig. 7).

Collectively, these results suggest that VcpR positively regulates transcription and secretion of T6SS1. Conversely, although VcpR positively regulates transcription of T6SS2 at a low level, functional secretion of Hcp2 is negatively controlled by QS, leading to highest secretion of Hcp2 in LCD-mimicking mutants.

### QS activation of T6SS1 via VcpR is required for interbacterial competition

To determine the physiological effect of QS signaling on T6SS function, we focused on the ability of T6SS1 to mediate interbacterial competition. To this end, we competed *Vcor* isogenic strains against *V. natriegens*, a prey strain that does not possess a T6SS. In agreement with the results observed in the VgrG1 secretion assay in Figure 7, T6SS1-mediated killing of prey bacteria was abrogated in the Δ*vcpR* mutant as shown previously (36) and in the Δ*luxP* mutant (Fig. 8). These results were similar to the effect of T6SS1 inactivation by *hcp1* deletion (Fig. 8). Notably, the prey bacteria grew more when competed against the Δ*vcpR* and Δ*luxP* mutant attacker strains than when competed against the Δ*hcp1* strain during the 4 hour-long incubation, suggesting that QS regulates antibacterial determinants other than T6SS1.

**Figure 8.**
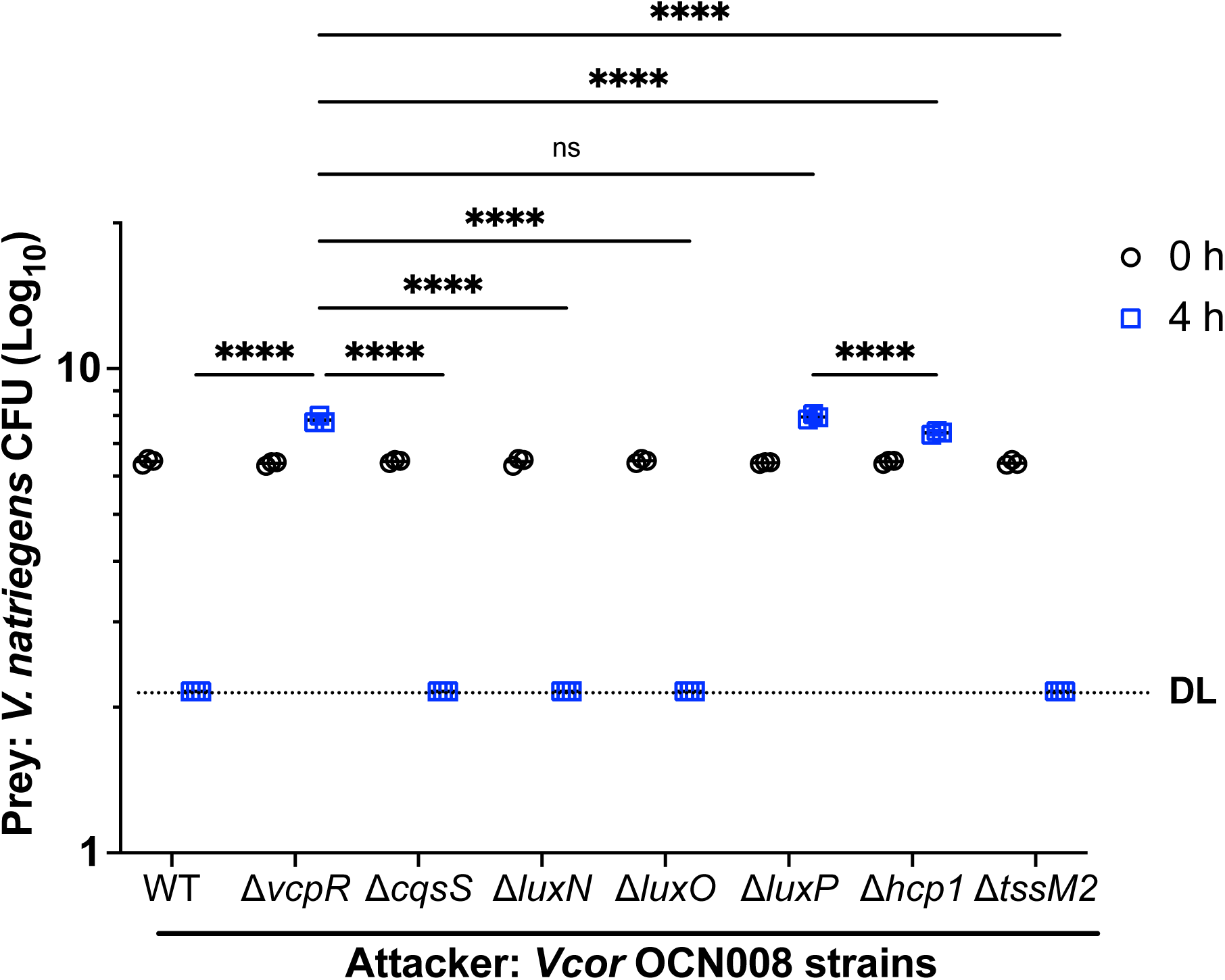
*Vcor* OCN008 outcompetes *V. natriegens* using T6SS1 and other potential virulence factors regulated by VcpR. Viability counts (colony forming units: CFU; Log_10_ transformed data) of *V. natriegens* prey strains before (0 h) and after (4 h) co-incubation with the indicated *Vcor* OCN008 attacker strains. Asterisks indicate significance (two-way analysis of variance; **** = *p*<0.0001, ns = not significant (*p* > 0.05); *n* = 3 for each assay) on normally distributed data (D’Agostino & Pearson test) followed by Šídák’s multiple comparisons test, calculated with log-transformed data. WT, wild-type; DL, assay detection limit.

Further, single deletions of either *cqsS* or *luxN* were not sufficient to alter killing; the toxicity of these strains was not significantly different from wild-type (Fig. 8). From these data, we conclude that QS signaling regulates T6SS1-mediated killing through VcpR-dependent transcriptional activation of this system.

## DISCUSSION

A considerable number of studies implicate *Vcor* as a causative agent for coral diseases and, in specific coral species, a non-virulent, commensal microbe (8, 9, 58, 89). However, the mechanistic processes regulating *Vcor* pathogenicity and virulence gene expression are not clearly defined and likely vary among hosts. QS is a prominent regulatory process for pathogenic behaviors in *Vibrio*-infected marine hosts (20, 37, 90–92). Previous metagenomic studies identified QS as a process, among many others, likely involved in coral bleaching (18, 28, 66). In this study, we have established that three species of *Vcor* encode at least one active QS signaling pathway that controls the QS master regulator VcpR. Specifically, in OCN008, VcpR controls the expression and function of at least three potential virulence factors: biofilm formation, protease activity, and T6SS. We postulate that VcpR also controls these native processes in all *Vcor* strains, and that virulence factors regulated by QS are involved in coral disease progression.

To define the genes controlled by QS in *Vcor*, we examined the regulon of VcpR in two strains: OCN008 and BAA-450. As a comparison, the VcpR homologue in *V. campbellii*, LuxR, activates and represses hundreds of genes responsible for regulating colonization and virulence behaviors (30, 42, 43, 93, 94). Likewise, we observed that VcpR regulons in two strains include T6SS1 genes, proteases, biofilm genes, and more, aligning with the regulons of LuxR/HapR proteins in other *Vibrio* spp. The size of the regulons of VcpR in OCN008 and BAA-450 differed substantially: 896 versus 363, respectively. The observed difference in QS regulon size between strains also aligns with the examination of QS regulons in strains of *V. campbellii* where the BB120 strain has a larger QS regulon than DS40M4 (42). These lists of genes will support future studies of how QS affects virulence mechanisms.

Although we could easily detect the production of DPD (which forms AI-2) in all three *Vcor* isolates, we failed to detect AI-1 or CAI-1. It is not surprising that we did not identify AI-1 given that each *Vcor* isolate encodes a gene product with weak homology to *V. campbellii* LuxM, and this gene precedes either 1 or 2 copies of a LuxN homolog. The conservation of these genes is generally low comparing *Vcor* to other *Vibrio* species; for example, *Vcor* OCN008 LuxM shares 43% identity with *V. campbellii* LuxM, and *Vcor* OCN008 LuxN1/N2 share 43% identity with *V. campbellii* LuxN (Fig. 2, S1). However, there is a wide range of conservation across *Vcor* strains for these two genes; some strains have nearly identical proteins to OCN008 LuxM/LuxN (98-100%), whereas others are much more divergent (23-74%). Further, in some *Vcor* strains, there were duplicate *luxN* genes, though these paralogs share only 46% identity with each other in OCN008 (Fig. 2B). Although these genes are maintained in *Vcor* strains, their sequence diversity suggests that these enzymes have evolved to produce and detect different signals. Further studies of the activities of the LuxM/LuxN gene products in *Vcor* species will enable us to understand if they are signaling and identify the structure of the signaling molecule(s).

Conversely, the lack of detection of CAI-1 was intriguing; *Vcor* strains encode homologs with ∼59% identity to *V. campbellii* CqsA (Fig. 2), and likewise, the CqsS sensor was moderately conserved at ∼49%. We postulate that these organisms may make a CAI-1-like molecule, but it may contain a variation that has prevented its detection thus far. For example, *V. cholerae* produces CAI-1, whereas *V. campbellii* produces enamino-CAI-1 (86, 95, 96). Notably, the CqsA homologs in *V. cholerae* and *V campbellii* share only 58% identity. Although both enzymes use CoA-hydrocarbons and *S-*adenosylmethionine (SAM) as the substrates, the mechanism for the substrate specificity (length and type of hydrocarbon) that dictates the production of CAI-1 (C10-CAI-1) versus Ea-C8-CAI-1 is not understood. Thus, it is possible that the *Vcor* CqsA also produces a modified CAI-1-like molecule, and we hypothesize that *Vcor* CqsS has evolved to sense that particular molecule. Analyses of the production profile of CqsA from *Vcor* in future studies will likely reveal the molecule produced by *Vcor* strains. In summary, we determined LuxN1 and LuxN2 do not produce a functional signaling molecule, similar to AI-1, nor could we conclude the production and detection of CAI-1 by CqsA and CqsS, respectively. More importantly, AI-2 is synthesized in *Vcor* and sensed by LuxPQ thus completing the signaling pathway to further regulate QS phenotypes.

We found that the T6SS1 and T6SS2 main gene clusters are differentially regulated at the transcriptional level by QS; most T6SS1 core genes are highly induced by VcpR, whereas only a few of the T6SS2 core genes are induced above the 2-fold cut-off in the RNA-seq analysis. Surprisingly, the effect of QS on T6SS1 and T6SS2 activity is inversed. QS is required for activating T6SS1-mediated secretion, in agreement with the observed transcriptional regulation. Conversely, QS signaling activates transcription of several T6SS2 genes, yet T6SS2 secretion is induced in the Δ*luxP* and Δ*vcpR* strains that mimic a LCD-state. Moreover, the secretion of Hcp2 was absent in the Δ*luxO* strain, which mimics the HCD-state. Taken together, both results align to suggest that while VcpR transcriptionally regulates some genes in T6SS2, another mechanism – likely posttranscriptional – is controlling the activation, assembly, and firing of the T6SS2 apparatus.

Corals harbor a diverse set of innate microbes and subsequent challenges for invading *Vcor* cells to overcome and ultimately cause disease. To proliferate in that environment, *Vcor* likely utilizes an arsenal of virulence factors (59, 97, 98). Conditions such as inoculum dose, seawater temperature, and host cell interaction induce different infection states and gene regulation among *Vcor* strains (21, 27, 59, 66, 99). QS-regulated processes, like biofilm formation, protease production, and T6SS-mediated effector delivery, in addition to other behaviors previously reported in *Vcor*, collectively comprise virulence strategies aiding *Vcor* in host colonization, proliferation, circumventing host-immune response, and dissemination (36, 100–102). In conclusion, we identified an active QS signaling pathway and the QS-mediated expression of virulence factors (biofilm formation, protease production, and interbacterial competition via the T6SS) in the coral pathogen, *Vcor*. Collectively, these findings provide relevant information to the field of marine pathogens and coral disease, as well as potential strategies for preventing *Vcor* pathogenicity in the future.

## MATERIALS AND METHODS

### Strain collection and media types

A list of bacterial strains used in this study is organized in Tables S1. All *Vibrio* strains were grown at 30°C in LB marine (LM) medium (Lysogeny broth supplemented with 10 g NaCl L^-1^ for a total of 20 g NaCl L^-1^) and, for select bioluminescence assays, in M9 minimal salts supplemented with glucose (20 mM) and casaminoacids (0.2%) (M9GC) dissolved in purified water. Interbacterial competition assays and T6SS assays were performed in MLB, which is LB containing 3% NaCl. *Escherichia coli* transformants were grown in Super Optimal broth with Catabolite repression (SOC) (0.5% Yeast Extract, 2% Tryptone, 10 mM NaCl, 2.5 mM KCl, 10 mM MgCl_2_, and 10 mM MgSO_4_) and supplemented with filter sterilized glucose to a final concentration of 20 mM after medium sterilization. In instances of transformations and conjugations involving an *E. coli* diaminopimelic acid (DAP) auxotroph strain (*dapA*), the media were supplemented with 0.3 mM DAP. Ex-conjugations between the recipient *Vcor* strain and donor *E. coli* plasmid strain were plated on LB plates, and recipient exconjugants were selected on LM plates with selective antibiotics and DAP. Antibiotic concentrations used in *Vcor* and *E. coli* were kanamycin (100 µg mL^-1^ in LM and 40 µg mL^-1^ in LB media), chloramphenicol (10 µg mL^-1^), and gentamicin (100 µg mL^-1^).

### Gene knockout plasmid construction

The plasmids and DNA oligonucleotides used in this study are listed in Tables S2 and S3, respectively. The *E. coli* strain b914::pSW4426T (103) is an empty suicide vector used to create clean deletion plasmids for *Vcor*. To construct *Vcor* mutant strains, ∼1000 base pairs upstream (up) and downstream (down) of the gene (*e.g., vcpR* and *luxO*) were PCR-amplified using wild-type (WT) *Vcor* genomic DNA from strains BAA-450, OCN008, or OCN014 (Thermo Scientific GeneJET Genomic DNA Purification Kit) and cloned into plasmid pSW4426T as described (27). Cloning procedures used oligos listed in Table S3 and methods are available upon request. *E. coli* strain b3914 was used for all cloning procedures and selected with DAP and chloramphenicol. Plasmids were sequenced at Eurofins Scientific. Additionally, samples underwent centrifugation using Eppendorf microcentrifuge model 5424 and rotor FA-24×2.

For conjugation of knockout plasmids from *E. coli* to *Vcor*, donor and recipient strains were grown overnight in their representative LB media, antibiotics, and/or DAP as required. Before conjugating, *E. coli* and *Vcor* strains grown with antibiotics were washed by pelleting 1 mL of overnight culture at 15,871 xg, discarding the supernatant, and resuspending the cells in LB or LM respectively, and DAP, when required. Once washed, the *E. coli* donor plasmids and *Vcor* recipient strain were combined and mixed in a mating spot on a LB agar plate and incubated at 30°C for a minimum of four hours. To improve conjugation efficiency, some crossover matings included the helper plasmid pRK600, which facilitates increased pili production and aids in mobilizing the donor plasmid. The mating spot was streaked on an LM agar plate containing antibiotic for selection and grown at 30°C to select for antibiotic-resistant colonies. Ex-conjugants were screened via colony PCR or PCR using extracted genomic DNA and appropriate primers (Table S3) to confirm single-crossover recombination at the target locus. Strains with confirmed plasmid integration were next counter-selected for a second crossover recombination by inducing the *ccdB* toxin cassette via arabinose induction on 0.3% arabinose agar plates for 24-48 hours at 30°C. The plating process on LM and arabinose was repeated twice to ensure counter-selection. Deletion strains were confirmed by PCR and sequencing at the target locus.

### Bioluminescence assays

In preparation for bioluminescence curve assays, overnight bacterial cultures were grown in LM media and back-diluted 1:10,000 in 270 µL of fresh M9GC media and, when necessary, antibiotics. Samples were organized in a 96 black-welled, clear-bottom plate, skipping wells and rows between samples to minimize light carryover. OD_600_ and bioluminescence (gain set to 160) measurements were taken every 30 minutes for 19-22 hours using a BioTek Cytation 3 Plate Reader set to an internal temperature of 30°C and shaking between reads.

For the modified bioluminescent curve assays, overnight *Vcor* OCN008 containing the *luxCDABE* reporter plasmid (pCS18) was back-diluted 1:200 in LM and kanamycin and grown to OD_600_ = 0.1. The overnight test strains, *V. campbellii* BB120 (=ATCC BAA-1116) and *Vcor* OCN008, OCN014, and BAA-450, were back-diluted 1:10 in LM and grown to ∼ OD_600_ = 1.5; 1 mL of each strain was pelleted at 15,871 xg and 500 µL of each supernatant was filtered in a centrifugal filter tube (0.22 µM filter size). After all strains reached their target OD_600_, 160 µL of OCN008::pCS18, 40 µL of fresh media (as a control) or filtered supernatant, and 10 µL of boric acid (5 mM) were combined per well. Sample wells were organized in the plate and grown according to the above instructions. Measurements were taken every 30 minutes for 4 hours.

The autoinducer reporter assays in this study are modified from the method described by Simpson *et al*. (42). All overnight strains, including BB120 type-strains and *Vcor* strains intended for supernatant collection, were back-diluted 1:100 in fresh media and grew shaking at 30°C to OD_600_ = 1.5. Supernatant from the *Vcor* strains and BB120 (control) were collected and filtered in a centrifugal filter tube. Each filtered supernatant was added to fresh media (1:4 ratio) per well in a 96 well cell culture plate. At a final back-dilution of 1:5000, BB120 type-strains were added to the wells containing each *Vcor* supernatant-fresh media mixture and further supplemented with boric acid (5 mM). Control wells contained fresh media, back-diluted BB120 type strain cell culture, no additive supernatant, and in one set, boric acid. The 96 well cell culture plate was sealed with microporous tape to minimize evaporation, incubated at 30°C, and shaking at 275 RPM with lid covered for 18 hours. Once grown, samples from each well were transferred to a 96 black-welled, clear-bottom plate, skipping wells and rows between samples. OD_600_ and bioluminescence (gain set to 160) were measured using a BioTek Cytation 3 Plate Reader.

### Bioinformatics analysis of bacterial genomes

The heat-map table data was generated by Indiana University’s Center for Genomics and Bioinformatics in Bloomington, IN. Protein sequence sets for the *Vibrio campbellii* BB120 and the 14 *Vcor* strains were combined into a single sequence set which was clustered using cd-hit ver. v4.8.1-2019-0228 (42, 104) with the minimum sequence identity cutoff set at 40% (parameters: -n 2 -T 0 -M 0 -g 1 -s 0.8 -c 0.40). The clusters associated with the quorum sensing genes from BB120 were obtained, and the protein sequences for the top hits from each species within each cluster were extracted and tested with multiple sequence alignment (Fig. S1). A similar analysis was completed using the OCN008 genome sequenced by our group (63). All the OCN008 quorum sensing genes with at least 40% sequence identity to their corresponding BB120 homologs were identified from the corresponding clusters, except for *luxM* and *vpsS* genes. For those two genes, profile HMMs were built from the multiple sequence alignments of the corresponding homologs from other *Vcor* strains and searched against the *Vibrio* sequence set using hmmer ver. 3.2.1 (http://hmmer.org). The OCN008 gene G3U99_RS10200 was found to be a potential luxM homolog at 30% identity to BB120. For the OCN008 quorum sensing genes, thus identified, percent identity values to the corresponding potential homologs from other *Vcor* strains and BB120 were computed using BLASTP picking the best scoring hit.

### Biofilm formation assays and protease activity assays

Biofilm assays with *Vcor* OCN008 strains were performed as described (42). The protease activity assays in this study were completed using the nonspecific protease substrate dye, azocasein. Growth conditions and back-dilution method were completed as previously described (42), however, the back-diluted strains were grown overnight and culture OD_600_ was measured in a spectrophotometer. For cell collection, 1 mL of each culture was centrifuged at 6010 xg for 10 minutes, and 100 µL of supernatant was incubated with 400 µL of 1% azocasein for 30 minutes in a 37°C standing incubator. The reaction was stopped by adding 600 µL of 10% trichloroacetic acid (TCA), incubated on ice for 30 minutes, and centrifuged at 11,600 xg for 5 minutes. A cuvette containing 200 µL of 1.8 1N NaOH was mixed with 800 µL of supernatant. Using LM as a calibration blank, total protease activity was measured at OD_420_. Protease activity for each strain was normalized by dividing OD_420_ by OD_600_.

### Mass spectrometry of bacterial supernatants

Supernatant extracts from wild-type *Vcor* OCN008, OCN014, and BAA-450 and controls *V. campbellii* BB120 and *V. cholerae* C6706 were analyzed via mass spectrometry to identify autoinducer molecules or derivatives of AI-1, AI-2, and CAI-1.

**AI-1:** Supernatants for AI-1 detection were collected and extracted analogously to previous work (42) with the initial modification of mixing 50 mL of extracted supernatant with dichloromethane (3 x 20 mL) (Fig. S2).

**AI-2/DPD:** Separate supernatant preparation was performed to differentiate the analyses between AI-2 and DPDQ, a derivative of the AI-2 precursor molecule, DPD. For context, the autoinducer synthase protein, LuxS, produces the metabolic byproduct DPD, 4,5-Dihydroxy-2,3-pentanedione, which enzymatically interacts with boric acid to form AI-2, a furanosyl borate diester (105). For analysis, cultures were supplemented with 100 mM boric acid, which activated a conformational change of DPD into AI-2 (Fig. S3A), and 250 µL of supernatant were collected for AI-2 mass spectrometry analyses. Similarly, 250 µL of supernatant (without the addition of boric acid) was supplemented with 3 µL of 1 mM orthophenylene damine (OPD) to induce a conformational change of DPD molecules into DPDQ (Fig. S3B), and samples were incubated at room temperature for one hour prior to DPDQ detection. One microliter of sample solution was injected into the LC-MS system. The system consisted of an Agilent 1100 series LC system with a binary pump, thermostatted column compartment, and autosampler utilizing a Waters SymmetryShield C18 column 4.6 x 150 mm with 3.5 µm particles. Mobile phase A was water with 0.1% formic acid and mobile phase B was acetonitrile with 0.1% formic acid. The gradient consisted of a linear change from 0% B initially to 100% B over 10 minutes before being held at 100% B for 1 minute and then returning to start conditions over 1 minute. The total run time lasted 16 minutes. The mass spectrometer detector is an LTQ Orbitrap XLD system operated in positive electrospray ionization mode for this measurement. DPDQ is identified by comparison to a previously run standard and confirmed via high resolution mass analysis.

**CAI-1:** Two supernatant sample sets, derivatized and underivatized, were prepared for CAI-1 analysis. Though the initial report of CAI-1 (95) indicated that it could be detected via GC-MS without derivatization, a derivatization protocol was followed in an attempt to increase sensitivity for measurement of the molecule. For derivatized samples, filtered supernatants were first vacuum concentrated in a Speedvac without heat for 16-18 hours and then resuspended in 133 µL of methoxyamine hydrochloride (20 mg mL^-1^ in pyridine) and incubated for 30 minutes at 80°C. Each mixture received 177 µL of bis(trimethylsilyl)trifluoroacetamide (BSTFA) reagent [1% trimethylchlorosilane (v/v)] and was incubated for 1.5 hours at 70°C. To enhance the detection of underivatized CAI-1, we increased the final volume (500 mL) of each strain’s filtered supernatant. The samples were dried and then resuspended in dichloromethane before injection into the machine. Additionally, a CAI-1 standard ((3S)-3-Hydroxy-4-tridecanone) was purchased from Toronto Research Chemicals (cat. #C431460) and used for confirmation during mass spectrometry analyses.

Samples were analyzed using an Agilent 7890B/7250 GC-QToF system with a 20m DB-5MS column, also from Agilent. Using the purchased standard, we were able to confirm that the instrument was sensitive to the CAI-1 molecule to at least 2.5 µg mL^-1^ concentration when dissolved in dichloromethane. Because no strains examined in this study showed signal from this form of CAI-1, we used the LTQ Orbitrap XL instrument mentioned above in positive atmospheric chemical ionization (APCI) mode to look for other forms and were able to detect a small signal consistent with an enamine form of CAI-1 by a flow injection approach.

### RNA extraction, quantitative reverse transcriptase real-time PCR, and RNA-sequencing

Cell growth conditions, collection at OD_600_=1.5, RNA extraction, and reverse transcription quantitative PCR (RT-qPCR) were performed as described (42). RNA extracts were purified using QIAGEN RNeasy Micro Kit prior to RNA-sequencing (RNA-seq). RNA-seq was performed and analyzed as described (106). The RNA-seq raw data were deposited in NCBI GEO: accession GSE232809.

### T6SS secretion assays

*Vcor* strains were grown overnight in appropriate media, then back-diluted 1:4 in fresh media and incubated for 2 additional hours at 28°C. Samples were then normalized to an OD_600_=0.18 in 5LJmL of MLB broth, and bacterial cultures were incubated with constant shaking (220LJrpm) at 28°C for 3.5 hours. For expression fractions (cells), cells equivalent to 0.5 OD_600_ units were harvested, and cell pellets were resuspended in 35 µL of 2x protein sample buffer (Novex, Life Sciences) with 5% (v/v) β-mercaptoethanol. For secretion fractions (media), supernatant volumes equivalent to 5.0 OD_600_ units were filtered (0.22 µm), and proteins were precipitated using the deoxycholate and trichloroacetic acid method (107). The precipitated proteins were washed twice with cold acetone and then air-dried before resuspension in 20 µL of 10 mM Tris-Cl (pH = 8.0) and 20 µL of 2x protein sample buffer with 5% (v/v) β-mercaptoethanol. Next, samples were incubated at 95°C for 10LJminutes and then resolved on a TGX Stain-free gel (Bio-Rad). The proteins were transferred onto 0.2 µm nitrocellulose membranes using Trans-Blot Turbo Transfer (Bio-Rad) according to the manufacturer’s protocol. Membranes were then immunoblotted with Custom-made α-VgrG1 (108) and α-Hcp2 antibodies (polyclonal; raised in rabbits against the peptide CVMTKPNREGSGADP; GenScript) (87), and Direct-Blot^TM^ HRP anti-*E. coli* RNA polymerase sigma 70 (mouse mAb #663205; BioLegend; referred to as α-RNAp) at 1:1000 dilutions. Protein signals were visualized in a Fusion FX6 imaging system (Vilber Lourmat) using ECL reagents. The assay was performed three times with similar results. A representative result is shown.

### Bacterial competition assays

Attacker and prey *Vibrio* strains were grown overnight in MLB broth. Attacker strains were back-diluted 1:10 into fresh media and incubated for an additional hour at 28°C. Attacker and prey cultures were then normalized to an OD_600_ of 0.5 and mixed at a 4∶1 (attacker: prey) ratio in triplicate. Next, 25 μL of the mixtures were spotted onto MLB agar competition plates and incubated at 28°C for 4LJhours. The colony-forming units (CFU) of the prey strains at timeLJ=LJ0LJhours were determined by plating 10-fold serial dilutions on selective media plates. After 4LJhours of co-incubation of the attacker and prey mixtures on the competition plates, the bacteria were harvested and the CFUs of the surviving prey strains were determined by plating 10-fold serial dilutions on selective media plates. *V. natriegens* prey strain contained a pBAD33.1 plasmid to allow selective growth on plates containing chloramphenicol. The assay was performed three times with similar results. A representative result is shown.

## ACKNOWLEDGEMENTS

The authors would like to thank members of the van Kessel, Salomon, and Ushijima labs for helpful discussions of this work and for comments on the manuscript. In addition, we thank Chelsea Simpson for advice and support in experimental design and interpretation. This project was funded by the National Science Foundation and US-Israel Binational Science Foundation (NSF-BSF) under award numbers IOS-2207168 (JCVK), IOS-2207169 (BU), and 2021733 (DS).

