## Supplemental Information for "Quorum Sensing Regulates Virulence Factors in the Coral Pathogen *Vibrio coralliilyticus*"

Supplemental Figures 1-4
Supplemental Tables 1-3

Figure S1

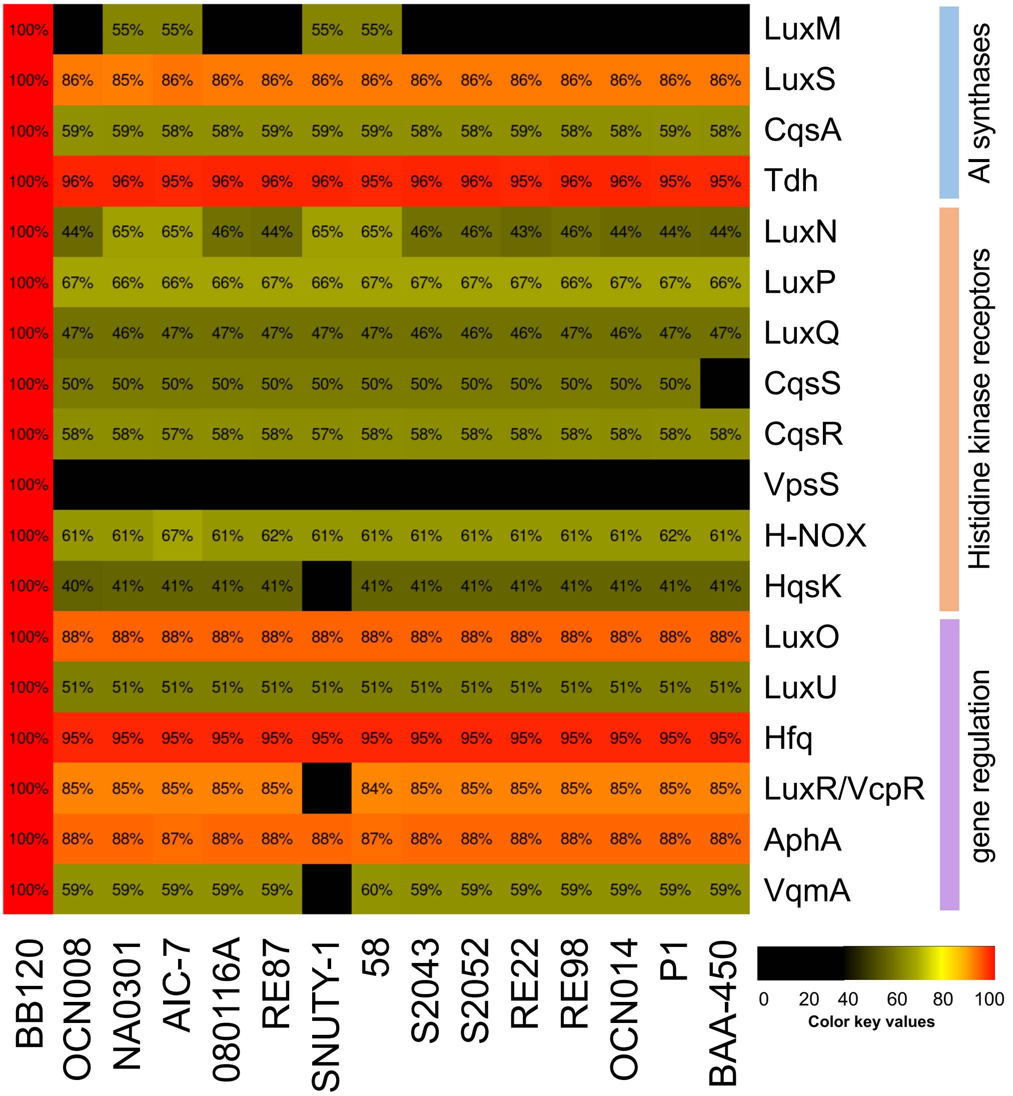


**Figure S1. Conservation of QS system homologs in *Vcor* strains.** The proteins listed on the y-axis are involved in QS in *V. campbellii* BB120 (query sequences) and are grouped based on function. The heat-map indicates conservation of these proteins in *Vcor* strains (amino acid identity); 40% identity or lower is colored in black.

Figure S2

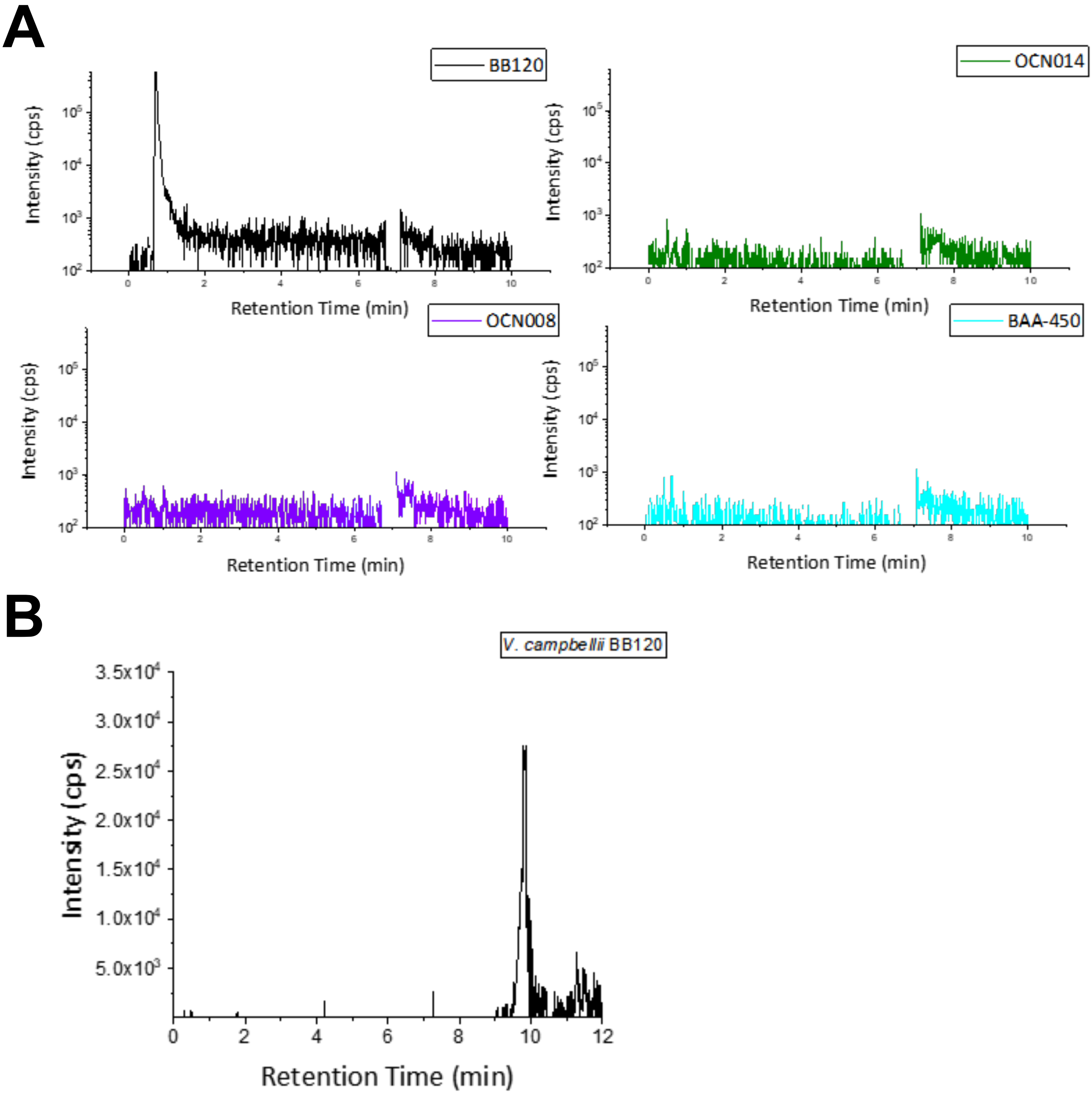


**Figure S2. Detection of autoinducers using mass spectrometry.** (A) Total ion chromatograms of AHL molecules in extracts of supernatant from strains *Vcor* OCN008, OCN014, and BAA-450, and positive control strain *V. campbellii* BB120. (B) Base peak chromatogram (BPC) and extracted ion chromatogram (EIC) detection of enamine CAI-1 from *V. campbellii* BB120 supernatant extract.

Figure S3

# A

### AI-2 Formation:

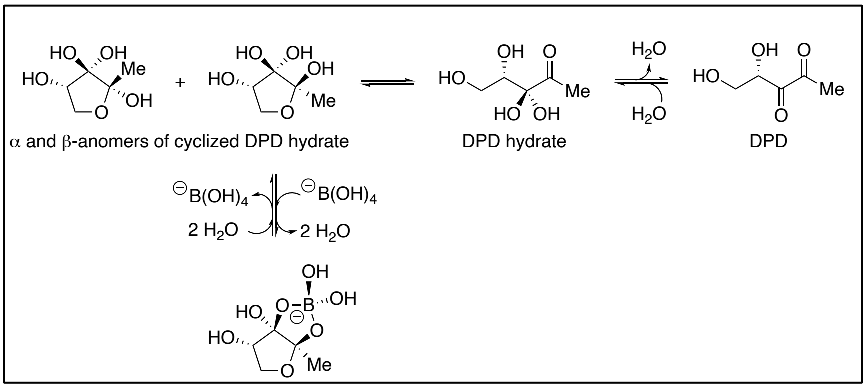


# B

### Derivatization and detection of DPDQ:

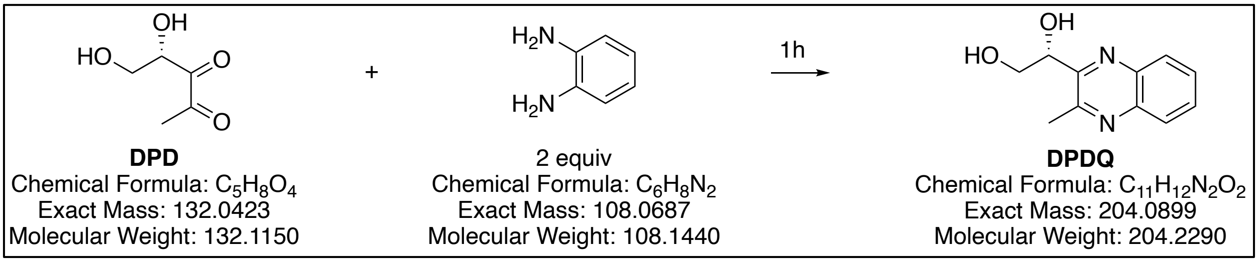


**Figure S3. AI-2 and derivatize compounds detected by mass spectrometry.** (A) AI-2 precursor molecule (DPD) and addition of boric acid to form AI-2. (B) DPD with the addition of chemical OPD forms the detectable compound, DPDQ.

Figure S4

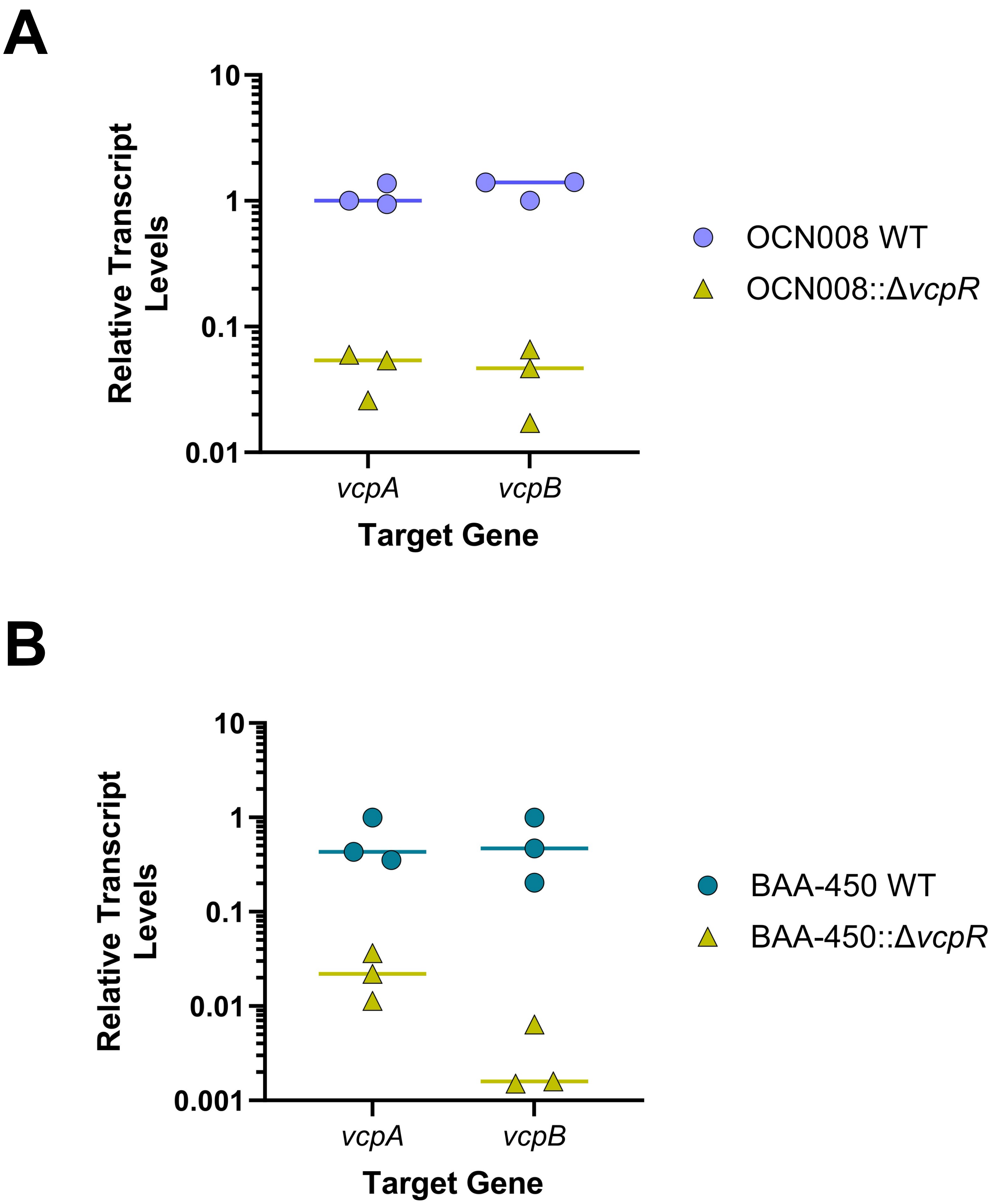


**Figure S4. VcpR regulates the protease-encoding genes, *vcpA* and *vcpB*.** Reverse transcriptase quantitative PCR (RT-qPCR) of transcript levels for target genes *vcpA* and *vcpB* comparing wild-type and Δ*vcpR Vcor* strains of parent OCN008 (A) and BAA-450 (B). *recA* was used as the standard control for analyses.

**Table S1. Bacterial strains used in this study.**

| Strains | | |
|---|---|---|
| **Name** | **Description** | **Reference** |
| OCN008 (VL008) | *V. coralliilyticus* type strain (wild type) | Ushijima et al. 2014 |
| BAA-450 (VL004) | *V. coralliilyticus* type strain (wild type) | American Type Culture Collection (ATCC), atcc.org |
| OCN014 (VL009) | *V. coralliilyticus* type strain (wild type) | Ushijima et al., 2016 |
| BB120 | *V. campbellii* type strain (wild type) | Bassler et al., 1997 |
| VL010 | OCN008::Δ*luxR* | Guillemette et al., 2020 |
| VL030 | OCN008::Δ*luxO* | Lab Collection |
| VL027 | OCN008::Δ*luxN* | Lab Collection |
| VL028 | OCN008::Δ*luxP* | Lab Collection |
| VL029 | OCN008::Δ*cqsS* | Lab Collection |
| VL034 | OCN008::pCS18, $kan^R$ | This study |
| VL035 | OCN008::Δ*vcpR*, pCS18, $kan^R$ | This study |
| VL044 | OCN008::pCS19, $kan^R$ | This study |
| VL045 | OCN008::Δ*vcpR*, pCS19, $kan^R$ | This study |
| VL018 | BAA-450::Δ*vcpR* | This study |
| VL019 | BAA-450::Δ*luxO* | This study |
| VL038 | BAA-450::pCS18, $kan^R$ | This study |
| VL011 | OCN014::Δ*vcpR* | This study |
| VL036 | OCN014::pCS18, $kan^R$ | This study |
| VL037 | OCN014::Δ*vcpR*, pCS18, $kan^R$ | This study |
| KM816 | BB120::Δ*luxS* | Waters and Bassler, 2006 |
| TL203 | BB120::Δ*cqsA* | Simpson et al., 2019 |
| JMH363 | BB120::Δ*luxM* | Waters and Bassler, 2006 |
| TL189 | BB120::Δ*luxS, ΔcqsA, ΔluxM* | Simpson and Petersen et al., 2020 |
| VL066 | OCN008:: Δ*hcp1* | Mass et al, 2024 |
| VL067 | OCN008:: Δ*tssM2* | Mass et al, 2024 |
| ecVL017 | *E. Coli* b3914 (wild-type) | Lab Collection |

| | | |
|---|---|---|
| Vnat::pBAD33 | *Vibrio natriegens* ATCC 14048 wild-type::pBAD33; $Cm^R$ | Mass et al, 2024 |

**Table S2. Plasmids used in this study.**

| Plasmids | | |
|---|---|---|
| **Name** | **Description** | **Reference** |
| pCS18 | $P_{luxCDABE}$; $kan^R$; derivative of pMMB67EH-tfoX-kanR | Chelsea Simpson |
| pCS19 | $P_{luxCDABE}$::*gfp*; $kan^R$; derivative of pMMB67EH-tfoX-kanR | Simpson et al., 2019 |
| pCS42 | $P_{luxCDABE}$::*gfp*; $gent^R$; derivative of pMMB67EH-tfoX-gentR | Chelsea Simpson |
| pVND01 | *E. Coli* b3814::pSW4426T; $Cm^R$, $Sp^R$, $Sm^R$, R6k oriV, PBAD-ccdB; DAP auxotroph; suicide vector to delete LuxO in *Vcor* ATCC BAA-450 | This study |
| pVND02 | *E. Coli* b3814::pSW4426T; $Cm^R$, $Sp^R$, $Sm^R$, R6k oriV, PBAD-ccdB; DAP auxotroph; suicide vector to delete VcpR in *Vcor* ATCC BAA-450 | This study |
| pBU247 | *E. Coli* b3814::pBU247; $Cm^R$; DAP auxotroph; suicide vector to delete Tssm1 in *Vcor* OCN008 | Lab Collection |
| pRK600 | Helper plasmid | |
| pSW4426T | *E. Coli* b3814::pSW4426T; $Cm^R$, $Sp^R$, $Sm^R$, R6k oriV, PBAD-ccdB; DAP auxotroph; empty suicide vector | Le Roux et al., 2007 |

**Table S3. Oligonucleotides used in this study.**

| DNA oligonucleotides | | | |
|---|---|---|---|
| **Primer** | **Description** | **Sequence** | **Source** |
| VNL019 | Forward amplify pSW4426T backbone (universal) | 5’-tcgcacgatatacaggattttgcc-3' | This study |
| VNL062 | Reverse amplify pSW4426T backbone (universal) | 3’-gcgaaggatctcttcagctcagtc-5' | This study |
| VNL058 | Amplify downstream forward insert (vcpR) | 5’-TATATAGAATTCgaaaactacttatcatgggcggt-3’ | This study |

| | | | |
|---|---|---|---|
| VNL044 | Amplify downstream reverse insert (vcpR) | 3’-TATATACTCGAGattaaccagtgtcatcgtaaagccg-5' | This study |
| VNL043 | Amplify upstream forward insert (vcpR) | 5’-TATATACTCGAGagttatatttccttgccaattgagtt-3' | This study |
| VNL055 | Amplify upstream reverse insert (vcpR) | 3’-TATATATCTAGAtggcaagctgcaagatttag-5' | This study |
| VNL062 | Sequencing detection primer for section between UP and DOWN arms (pVND02, vcpR) | 5’-AAGGCATCAAGTACGAAGTAGC-3' | This study |
| VNL066 | Gene locus upstream forward (vcpR BAA-450) | 5’-gagtgaacgcagcaatcgacaac -3’ | This study |
| VNL067 | Gene locus downstream reverse (vcpR BAA-450) | 3’-tgcgtaatgtgacggttgttaacg -5’ | This study |
| VNL046 | Amplify upstream forward insert (luxO) | 5’-TATATAgaattcaggacgaacgtgtgttggtcaccac-3’ | This study |
| VNL047 | Amplify upstream reverse insert (luxO) | 3’-TATATACTCGAGcaaaagatatttggacttttgtgtttgc-5’ | This study |
| VNL048 | Amplify downstream forward insert (luxO) | 5’-TATATACTCGAGTCGATCATGGAAGTGTTAAATCAAAC-3’ | This study |
| VNL057 | Amplify downstream reverse insert (luxO) | 3’-TATATAtctagaTTCGATGATGTTTGAATATCGCTTCA-3’ | This study |
| VNL064 | Gene locus upstream forward (luxO BAA-450) | 5’-aagttcactaacaacgtcagttggc-3’ | This study |
| VNL065 | Gene locus downstream reverse (luxO BAA-450) | 3’-gctgtgtcgctgagatatcgagc-5’ | This study |
| VNL063 | Sequencing detection primer for section between UP and DOWN arms (pVND01, luxO) | 5’-ccgtcgtagagagaaacaacaagc-3’ | This study |

| | | | |
|---|---|---|---|
| JCV612 | Forward sequence of PluxC PCR filter binding probe | 5’-cttatgaagtccatactttttcactgaa-3’ | Julia van Kessel |
| JCV613 | Reverse sequence of PluxC PCR filter binding probe | 3’-tttgcccatttattattaaaggtaagtgtttt-5’ | Julia van Kessel |
| VNL022 | qRT-PCR reference gene OCN008 recA forward | 5’-ATGAACAAATCGGAGAAAGTGATG-3’ | This study |
| VNL023 | qRT-PCR reference gene OCN008 and BAA-450 recA reverse | 5’-gcataatagagcctttaccgaattg-3’ | This study |
| VNL069 | qRT-PCR reference gene BAA-450 recA forward | 5’-ttgattgcctcagaggactc-3’ | This study |
| VNL024 | qRT-PCR OCN008 and BAA-450 vcpA forward | 5’-atggttaaagttgaaaacgatgc-3’ | This study |
| VNL025 | qRT-PCR OCN008 and BAA-450 vcpA reverse | 5’-ctcgtttcacttcagcgaa-3’ | This study |
| VNL026 | qRT-PCR OCN008 and BAA-450 vcpB forward | 5’-atgaaaatagccaagcgttttt-3’ | This study |
| VNL027 | qRT-PCR OCN008 and BAA-450 vcpB reverse | 5’-agttgattggtttgcttaaataaact-3’ | This study |
| VNL070 | Amplify gene locus OCN008 recA forward | 5’-atgaacaaatcggagaaagtgatgg-3’ | This study |
| VNL071 | Amplify gene locus OCN008 recA reverse | 5’-ttatagctcttcttgctctggcatt-3’ | This study |
| VNL072 | Amplify gene locus BAA-450 recA forward | 5’-ttgattgcctcagaggactctgg-3’ | This study |
| VNL073 | Amplify gene locus BAA-450 recA reverse | 5’-ttatagctcttcttgctctggcatt-3’ | This study |
